# Stress diminishes outcome but enhances response representations during instrumental learning

**DOI:** 10.1101/2021.02.12.430935

**Authors:** Jacqueline Katharina Meier, Bernhard P. Staresina, Lars Schwabe

## Abstract

Stress may shift behavioural control from a goal-directed system that encodes action-outcome relationships to a habit system that learns stimulus-response associations. Although this shift to habits is highly relevant for stress-related psychopathologies, limitations of existing behavioural paradigms hindered previous research to answer the fundamental question of whether the stress-induced bias to habits is due to impaired goal-directed or enhanced habitual processing (or both). Here, we leveraged EEG-based multivariate pattern analysis to decode neural outcome representations, crucial for goal-directed control, and response representations, essential for habitual responding, during instrumental learning. We show that stress reduces outcome representations but enhances response representations, both of which were directly associated with a behavioural index of habitual responding. Further, changes in outcome and response representations were uncorrelated, suggesting that these may reflect distinct processes. Our findings indicate that habit behaviour under stress is the result of both enhanced habitual and diminished goal-directed processing.

## Introduction

Adaptive behaviour in complex environments requires an intricate balance of deliberate action and efficient responding. For instance, in the supermarket, faced with numerous products, it may certainly be helpful to weigh the pros and cons of a specific product before making a choice, yet thinking for hours about which toothpaste to buy may interfere with the goal of being home before dinner. The balance of thorough deliberation and efficiency is supported by at least two systems of behavioural control that operate in tandem: (i) a goal-directed systems that learns when to perform which action in order to achieve a desired outcome, in the form of stimulus (S) – response (R) – outcome (O) associations and (ii) a habit system that acquires S-R associations, without any links to the outcome engendered by the response (***Adams, 1982; Adams and Dickinson, 1981; Balleine and Dickinson, 1998***). These systems are known to rely on distinct neural circuits. Whereas the goal-directed system is mainly based on the medial prefrontal and orbitofrontal cortex as well as on the dorsomedial striatum, the habit system is primarily dependent on the dorsolateral striatum *(**Balleine and Dickinson, 1998; Balleine and O’Doherty, 2010; Corbit and Balleine, 2003; Ostlund and Balleine, 2005; Tricomi et al., 2009; Valentin et al., 2007; Yin et al., 2006**).* Moreover, it is commonly assumed that the goal-directed system guides early learning, while the habit system takes over once a behaviour has been frequently repeated. Thus, buying toothpaste for the first time should be a goal-directed action, while this choice should become more habitual if we have bought a specific toothpaste several times before. Adaptive behaviour requires the capacity to flexibly switch back from habitual to goal-directed control when the environment changes (e.g., when the previously bought toothpaste is out of stock). Lack of this flexibility in behavioural control may be detrimental to mental health. In particular, the overreliance on habitual responding has been linked to several mental disorders, including drug addiction, obsessive-compulsive disorder, schizophrenia, eating disorders, and depression *(**Everitt and Robbins, 2005; Gillan et al., 2015; Griffiths et al., 2014; Robbins et al., 2012; Voon et al., 2015**).*

Accumulating evidence indicates that stressful events tip the balance from goal-directed to habitual control. Specifically, stress and major stress mediators have been shown to induce a shift from goal-directed action to habitual responding *(**Braun and Hauber, 2013; Dias-Ferreira et al., 2009; Gourley et al., 2012; Schwabe et al., 2012; Schwabe and Wolf, 2009, 2010, 2013; Smeets et al., 2019; Soares et al., 2012**).* Beyond its crucial relevance for our understanding of behavioural control in general, this stress-induced shift towards the habit system may be a driving force in mental disorders that are characterized by dysfunctional stress responses on the one hand and aberrant habitual control on the other hand *(**Adams et al., 2018; Goeders, 2004;Schwabe et al., 2011a**).* Although the stress-induced bias to habitual responding may have critical clinical implications, the exact mechanisms through which stress modulates the balance of goal-directed and habitual control are not fully understood.

In particular, previous research was unable to address the fundamental question whether the stress-induced bias towards habit behaviour is due to a down-regulation of the goal-directed system, an enhancement of the habit system, or both. Canonical assays for the assessment of goal-directed and habitual control do not allow a distinction between these alternatives. These paradigms are based on the key distinctive feature of goal-directed and habitual control, i.e. that goal-directed but not habitual control relies on the motivational value of the outcome (***Adams, 1982; Dickinson and Balleine, 1994***). Accordingly, classical paradigms tested the behavioural sensitivity to either a devaluation of an outcome or to the degradation of the action-outcome contingency (***Adams, 1982; Corbit and Balleine, 2003; Tanaka et al., 2008; Valentin et al., 2007; Yin et al., 2004***). Although these elegant paradigms provided valuable insights into mechanisms involved in behavioural control, they cannot show whether increased responding to devalued or degraded actions is due to reduced goal-directed or enhanced habitual control (or both).

Here, we aimed to overcome these shortcomings of classical paradigms for the assessment of the mode of behavioural control by leveraging electroencephalography (EEG) in combination with multivariate pattern analysis (MVPA)-based decoding of neural representations, in order to unravel whether acute stress enhances habitual processes, diminishes goal-directed processes, or both. In the present study, we exposed participants first to a stress or control manipulation and then asked them to complete an instrumental learning task during which they could learn S-R(-O) associations. Critically, we used image categories as response options (R) and outcome categories (O), respectively, that have a distinct neural signature and recorded EEG throughout the task. EEG-based classifiers (support vector machines, SVMs) were trained in a separate delayed-matching-to-sample task to distinguish between the R and O stimulus categories, respectively. We then applied these classifiers to the instrumental learning task to decode outcome representations, relevant for the goal-directed system, when participants saw the stimulus S and when they made the response R. Since the habit system builds strong S-R associations, we decoded response R representations when participants saw the stimulus S, assuming that habitual responding is reflected in enhanced R representations. We further included transient outcome devaluations after initial, moderate, and extended training in the instrumental learning task to assess if and how the predicted changes in neural representations are linked to behavioural manifestations of stress-induced changes in behavioural control.

## Results

The goal of this study was to elucidate the mechanisms underlying the impact of stress on the control of instrumental behaviour. Specifically, we aimed to leverage an EEG-based decoding approach to determine stress-induced changes in outcome and response representations that constitute key distinguishing features of goal-directed action and habitual responding *(**Adams, 1982; Adams and Dickinson, 1981; Balleine and Dickinson, 1998; Balleine and O’Doherty, 2010**).* To this end, participants first underwent the Trier Social Stress Test (TSST; ***Kirschbaum et al., 1993***), a mock job interview that represents a gold standard in experimental stress research *(**Allen et al., 2014**),* or a non-stressful control manipulation. Thereafter, participants completed a reinforcement learning task *(**Luque et al., 2017**)* that allowed us to probe the goal-directed vs. habitual control of behaviour. In this task, participants could learn S-R-O associations (Figure 1). Specifically, they acted as ‘space traders’ who traded two cards, represented by two distinct fractals (green vs. pink), with two alien tribes, represented by distinct symbols (red vs. blue), to receive cards from two possible categories (objects vs. scenes). On each trial, participants first saw one of the two fractals (S). They were then shown representatives of the two alien tribes next to each other and had to decide which alien to offer the fractal to (R). Finally, participants received feedback about whether the alien accepted the offer, traded one of the desired cards, and how many points were earned (O). Importantly, one alien tribe accepted only one type of fractal and traded only one card category. Furthermore, one card category was worth more than the other (high-valued outcome, O^high^ and low-valued outcome, O^low^). Participants had to learn these associations based on trial-by-trial feedback. Moreover, there was a response cost associated with each trade that was accepted by the alien.

**Figure 1.**
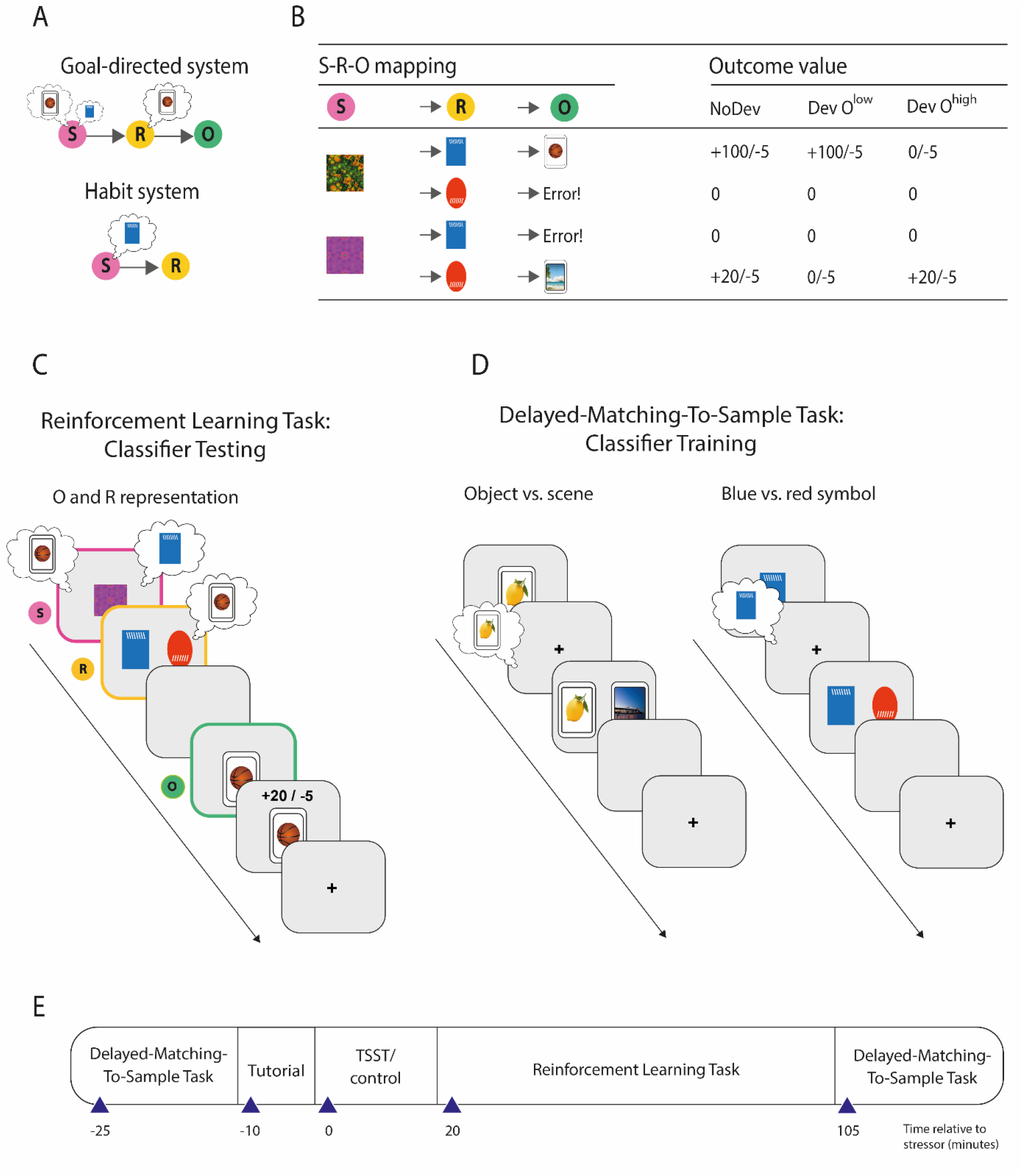
**A** Illustration of the goal-directed and habit system. While the goal-directed system encodes associations between stimulus (S), response (R), and outcome (O), the habit system acquires S-R associations, independent of the outcome engendered by the response. Accordingly, the goal-directed system relies on outcome representations, whereas the habit system does not. In contrast, response representations during stimulus presentations are assumed to be more pronounced during habitual processing. **B** S-R-O mappings in the reinforcement learning task and outcomes in trials, in which either none of the possible outcomes was devalued (NoDev), the outcome with lower value was devalued (Dev O^low^), or the outcome with the higher value was devalued (Dev O^high^). **C** Schematic of the reinforcement learning task, in which participants were trained on S-R-O sequences in a trial-by-trial manner. Using an EEG-based support vector machine (SVM), neural representations of the outcome stimuli (object vs. scene) were decoded during stimulus presentation and during response choice. Moreover, neural representations of the response options (blue vs. red alien) were decoded during stimulus presentation. **D** The SVM was trained in an unrelated delayed-matching-to-sample task (maintenance phase) that required participants to keep stimuli in mind that belonged to categories used as outcomes or response options during the reinforcement learning task. **E** Timeline of the experiment.

This task can be solved by ‘goal-directed’ action-outcome (S-R-O) learning and by ‘habitual’ stimulus-response (S-R) learning. In order to reveal the mode of control at the behavioural level, we presented task blocks in which one of the outcomes was devalued (i.e., not worth any points but associated with a response cost, thus resulting in a negative outcome). If behaviour is goal-directed, it should be sensitive to the outcome-devaluation and participants should avoid the devalued action. If behaviour is habitual, in turn, it should be less sensitive to the outcome devaluation and participants should be more prone to perform the frequently repeated, but now devalued response. To assess whether the balance of goal-directed and habitual behaviour and its modulation by stress depends on the extent of training, we presented devaluation blocks early during training, after moderate training, and again after extended training at the end of the task.

Critically, we recorded EEG during task performance and used stimulus categories as R and O stimuli that are known to have distinct neural signatures (***Bae and Luck, 2018; Cairney et al., 2018; Taghizadeh-Sarabi et al., 2015; Treder et al., 2020***). We trained EEG-based multivariate classifiers in an unrelated delayed-matching-to-sample (DMS) task to discriminate between stimulus categories that were used as S (blue vs. red symbols that differed also in shape and line orientation) and O (objects vs. scenes), respectively (Figure 1). The DMS task was completed both before and after the reinforcement learning task and the classifier was trained on the pooled trials of the two DMS sessions, thus ruling out any time-dependent biases of the classifiers. The trained classifiers were then applied to the reinforcement learning task to determine neural representations of R and O. Response representations were decoded during the S presentation, whereas O representations were decoded during both the S presentation and participants’ choice (R). Outcome representations at the time of S presentation and R are indicative for goal-directed control. Habitual control, on the other hand, should be indicated by increased R representations at the time of stimulus (S) presentation.

### Successful stress manipulation

Significant subjective and physiological changes in response to the TSST confirmed the successful stress induction. Compared to participants in the control group, participants exposed to the TSST experienced the treatment as significantly more stressful, difficult and unpleasant than those in the control condition (all *t*_56_ > 5.82, all *P* < 0.001, all *d* > 1.530, all 95% confidence intervals (CI) = 1.456 to 2.112; Table 1). At the physiological level, the exposure to the TSST elicited significant increases in pulse, systolic and diastolic blood pressure (time point of measurement × group interaction, all *F*(4, 224) > 13.55, all *P* < 0.001, all *η_p_^2^* > 0.195, all 95% CI = 0.100 to 0.425; Figure 2). As shown in Figure 2A-C, groups had comparable blood pressure and pulse before and after the TSST (all *t*_56_ < 2.28, all *P*_corr_ > 0.081, all *d* < 0.600, all 95% CI = 0.069 to 0.702), while participants in the stress group had significantly higher blood pressure and pulse than those in the control group during the experimental manipulation (all *t*_56_ > 4.21, all *P*_corr_ < 0.001, all *d* > 1.107, all 95% CI = 0.549 to 2.098). Finally, salivary cortisol concentrations increased in response to the TSST but not after the control manipulation (time point of measurement × group interaction: *F*(3, 168) = 6.69, *P* < 0.001, *η_p_^2^* = 0.107, 95% CI = 0.026 to 0.188). As shown in Figure 2D, participants of the stress and control groups had comparable cortisol concentrations at baseline (*t*_56_ = 1.16, *P*_corr_ = 1, *d* = 0.304, 95% CI = −0.216 to 0.821). However, about 20 minutes after the treatment, when the reinforcement learning task started, cortisol levels were significantly higher in the stress group than in the control group (*t*_56_ = 2.74, *Pcorr =* 0.032, *d* = 0.720, 95% CI = 0.185 to 1.249). As expected, cortisol levels returned to the level of the control group by the end of the task (60 minutes: *t*_56_ = 0.50, *P*_corr_ = 1, *d* = 0.130, 95% CI = −0.386 to 0.645; 105 minutes: *t*_56_ = 0.42, *P*_corr_ = 1, *d* = 0.111, 95% CI = – 0.405 to 0.625).

**Table 1.** Subjective responses to the TSST or control condition.

|  | Control |  | Stress |  |
| --- | --- | --- | --- | --- |
|  | M | SEM | M | SEM |
| *Subjective assessments* |  |  |  |  |
| Stressfulness | 16.79 | 3.45 | 62.33* | 4.44 |
| Unpleasantness | 15.36 | 3.51 | 57.00* | 5.90 |
| Difficulty | 14.29 | 3.35 | 51.33* | 5.29 |

Subjective assessments were rated on a scale from 0 ("not at all") to 10 ("very much"). \* $P < 0.001$ , Bonferroni-corrected, significant group difference.

**Figure 2.**
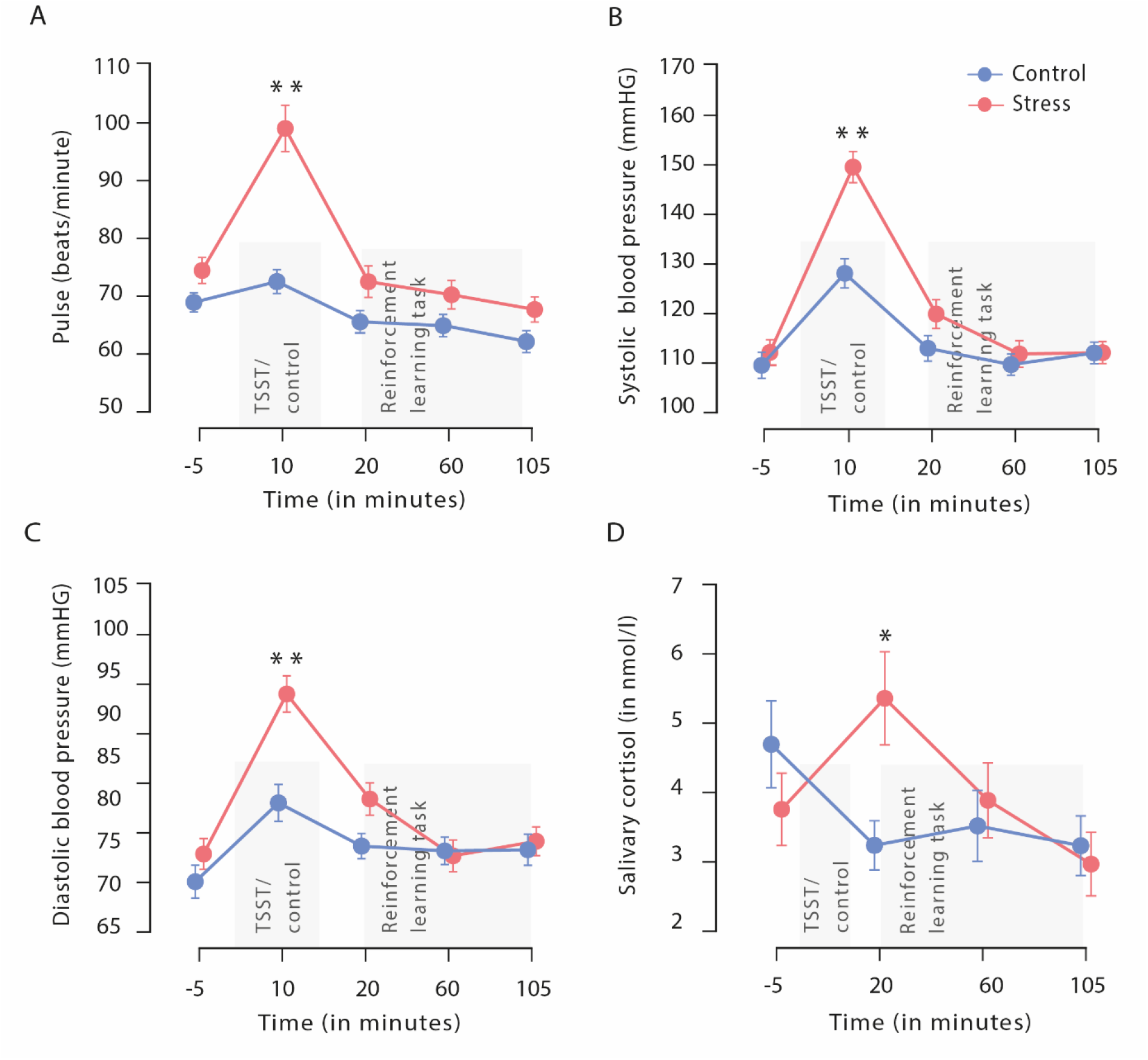
Physiological responses to the TSST. **A** Salivary cortisol (in nmol/L) increased significantly in response to the TSST but not in response to the control condition. Exposure to the TSST, but not to the control manipulation, resulted in a significant increase in systolic blood pressure (**A**), diastolic blood pressure (**B**), and pulse (**C**). The gray bars denote the timing and duration of the treatment (TSST vs. control condition) and the reinforcement learning task, respectively. Data represent means ± SEM. \**P* < 0.05, ** *P* < 0.001, Bonferroni-corrected, significant group difference.

### Stress renders behaviour less sensitive to outcome devaluation

Participants’ choice accuracy increased significantly across the task (*F*(2,114) = 10.08, *P* < 0.001, *η_p_^2^* = 0.150, 95% CI = 0.042 to 0.261) and reached an average performance of 94% correct responses in blocks without devaluation (NoDev), indicating that participants learned the task very well. During NoDev blocks, participants had a higher response accuracy for S^high^ than S^low^ trials (*t*_57_ = 3.29, *P* = 0.002, *d* = 0.432, 95% CI = 0.161 to 0.699, suggesting that learning was modulated by the value of the outcome. Performance in NoDev blocks was comparable in the stress and control groups (time × group interaction: *F*(2,112) = 2.44, *P* = 0.092, *η_p_^2^* = 0.042, 95% CI = 0 to 0.123; main effect group: *F*(1,56) = 0.30, *P* = 0.585, *η_p_^2^* = 0.005, 95% CI = 0 to 0.096), suggesting that stress did not affect instrumental learning per se.

During the reinforcement learning blocks, in which one of the outcomes was devalued (Dev), participants chose the action that was associated with the valued outcome significantly more often than the action that was associated with a devalued outcome (outcome devaluation × stimulus value interaction: *F*(2,110) = 163.31, *P* < 0.001, *η_p_^2^* = 0.785, 95% CI = 0.664 to 0.789; valued vs. devalued during Dev O^high^: *t*_57_ = 12.17, *P*_corr_ < 0.001, *d* = 1.589, 95% CI = 1.205 to 1.985; valued vs. devalued during Dev O^low^: *t*_56_ = 13.49, *P*_corr_ < 0.001, *d* = 1.786, 95% CI = 1.363 to 2.203), providing further evidence of successful instrumental learning. Importantly, there was also a significant outcome devaluation × stimulus value × time × group interaction *(F*(4, 220)= 4.86, *P* < 0.001, *η_p_^2^* = 0.081, 95% CI = 0.016 to 0.143). Follow-up ANOVAs revealed that stressed participants selected increasingly those actions that led to a devalued outcome during Dev O^high^ blocks at the end of the task (stimulus value × time × group interaction: *F*(2,112) = 8.89, *P* < 0.001, *η_p_^2^* = 0.137, 95% CI = 0.033 to 0.247; time × group interaction for devalued stimuli: *F*(2,112) = 9.09, *P* < 0.001, *η_p_^2^* = 0.140, 95% CI = 0.035 to 0.250; time × group interaction for valued stimuli: *F*(2,112) = 2.19, *P* = 0.116, *η_p_^2^* = 0.038, 95% CI = 0 to 0.116; block 2 vs. block 3 for devalued stimuli in the stress group: *t*_29_ = 4.28, *P*_corr_ = 0.001, *d* = 0.781, 95% CI = 0.366 to 1.186; block 2 vs. block 3 for devalued stimuli in the control group: *t*_27_ = 1.10, *P*_corr_ = 1, *d* = 0.207, 95% CI = −0.169 to 0.580), whereas there was no such effect during Dev O^low^ blocks (*F*(2,110) = 2.39, *P* = 0.097, *η_p_^2^* = 0.042, 95% CI = 0 to 0.124), nor during NoDev blocks (*F*(2,112) = 0.41, *P* = 0.667, *η_p_^2^* = 0.007, 95% CI = 0 to 0.052). As shown in Figure 3, stressed participants responded significantly more often to the devalued action than did non-stressed controls in the third devaluation block, at the end of the task (*t*_56_ = 2.61, *P*_corr_ = 0.036, *d* = 0.685, 95% CI = 0.152 to 1.213, stress vs. control during the first and second Dev O^high^ blocks: all *t*_56_ < 0.57, all *P*_corr_ = 1, all *d* < 0.149, all 95% CI = −0,472 to 0.664). These data suggest that stress rendered behaviour less sensitive to the outcome devaluation once the respective response has been intensively trained, which was in particular the case for the frequently repeated response to O^high^.

**Figure 3.**
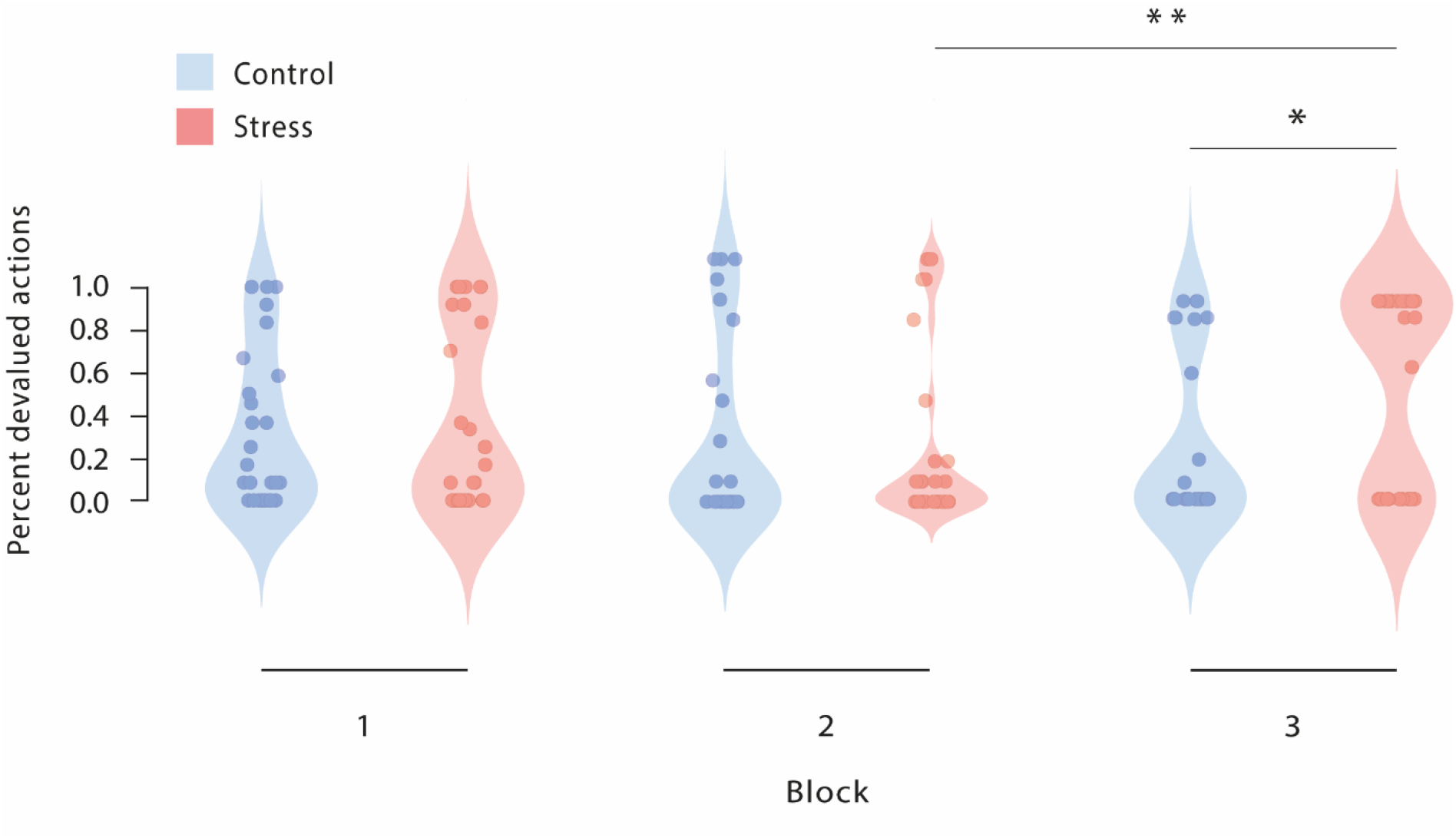
Percentage of responses to devalued actions across the reinforcement learning task during Dev O^high^ blocks. As training proceeded, stressed participants selected increasingly those actions that led to a devalued outcome. In addition, stressed participants responded significantly more often to the devalued action than non-stressed controls in the third devaluation block, at the end of the task. * *P* < 0.05, ** *P* < 0.01, Bonferroni-corrected. The data for Dev O^low^ and NoDev blocks are presented as supplemental material (Supplementary Figure 1 and Supplementary Figure 2, respectively).

Given that it is well-known that stress may disrupt memory retrieval (***Gagnon and Wagner, 2016; Quervain et al., 1998; Roozendaal, 2002***), it might be argued that the increased responding to the devalued action in stressed participants was due to their difficulty in remembering which outcome was valued and devalued, respectively. We could, however, rule out his alternative. Each block of our task involved, in addition to reinforcement learning trials, so called consumption trials in which participants could freely choose between the two outcome categories, without any response costs. These trials served to assess whether participants were aware of the current value of the outcomes. Here, participants clearly preferred the high-valued cards over low-valued cards during NoDev blocks (*F*(1,56) = 5382.91, *P* < 0.001, *η_p_^2^* = 0.990, 95% CI = 0.984 to 0.992) as well as the valued card over its devalued counterpart during Dev blocks (Dev O^high^ and Dev O^low^: both *F*(1,56) > 214.72, both *P* < 0.001, both *η_p_^2^* < 0.793, both 95% CI = 0.687 to 0.848; outcome devaluation × stimulus value interaction: *F*(2,112) = 876.42, *P* < 0.001, *η_p_^2^* = 0.940, 95% CI = 0.919 to 0.952), irrespective of stress (outcome devaluation × stimulus value × group interaction: *F*(2,112) = 0.05, *P* = 0.956, *η_p_^2^* = 0.001, 95% CI = 0 to 0.120). This finding demonstrates that participants of both groups were aware of the value of the card stimuli in a specific block but that responses of stressed participants were less guided by this knowledge about the value of the outcome engendered by the response.

Mean reaction times were significantly faster for high-valued stimuli than for low-valued stimuli during NoDev blocks (NoDev: *t*_57_ = 2.83, *P*_corr_ = 0.019, *d* = 0.372, 95% CI = 0.104 to 0.637), whereas participants responded faster to low-valued stimuli than to high-valued stimuli during Dev blocks (Dev O^low^: *t*_56_ = 5.73, *P*_corr_ < 0.001, *d* = 0.759, 95% CI = 0.462 to 1.052; Dev O^high^: *t*_57_ = 4.41, *P*_corr_ < 0.001, *d* = 0.579, 95% CI = 0.298 to 0.855; outcome devaluation × stimulus value interaction: *F*(2,110) = 28.14, *P* < 0.001, *η_p_^2^* = 0.338, 95% CI = 0.194 to 0.451). Stress did not influence participants’ reaction times (outcome devaluation × stimulus value × time × group: *F*(4,220) = 0.85, *P* = 0.494, *η_p_^2^* = 0.015, 95% CI = 0 to 0.043; main effect group: *F*(1,55) = 1.00, *P* = 0.322, *η_p_^2^* = 0.018, 95% CI = 0 to 0.134).

Although our analyses of the neural data focussed mainly on the decoding of outcome and response representations, there is recent evidence suggesting that habitual and goal-directed processes might also be reflected in event-related potentials (ERPs) (***Luque et al., 2017***). The extent to which a reward related ERP is sensitive to an outcome devaluation or not is assumed to indicate the degree of habitual or goal-directed processing. Thus, we additionally analysed stress effects on related ERP depending on outcome devaluation. Detailed information on these analyses are given in the supplemental information. In brief, we obtained that the occipital stimulus-locked P1 component was insensitive to outcome devaluation, suggesting the formation of an “attentional habit” *(**Luque et al., 2017***). This component was not modulated by stress. In contrast, a late component was sensitive to the outcome devaluation during Dev O^high^ blocks in control participants but not in stressed participants (Supplementary Figures 3–5). This pattern of results suggests that stress interferes with a late ERP component that has been linked to goal-directed processing as it was rather sensitive to the value of an outcome, whereas an outcome-insensitive ERP was present in both groups. However, similar to the behavioural response to a devalued action, the stress-induced decrease in a rather ‘outcome-sensitive’ ERP component leaves the question open as to whether this stress effect is due to a decrease in the goal-directed system or to an increase in the habit system.

### Stress reduces outcome representations after extended training

So far, our behavioural data showed that stress rendered behaviour less sensitive to a change in the value of an outcome, which can be interpreted as decreased goal-directed or increased habitual behaviour. In addition, stress reduced electrophysiological late latency potentials that appeared to be sensitive to the value of an outcome *(**Luque et al., 2017**).*

In a next step, we leveraged an EEG-based decoding approach to address the primary objective of this study, i.e., to probe the effect of stress on neural outcome and response representations, thereby providing insight into the neural signatures of the goal-directed and habit system, respectively. We trained an MVPA classifier (support vector machine, SVM) based on an independent dataset (DMS task, for details see materials and methods) to discriminate between categories that were used during the reinforcement learning task as outcome (object card vs. scene card). Then, this classifier was used to assess changes in outcome representations trial-by-trial throughout the reinforcement learning task. Goal-directed control should be reflected in high accuracy of outcome category classification during the presentation of the stimulus (S) as well as during the response choice (R).

We first analysed outcome representations during the presentation of the fractal stimulus S, before participants had to make a choice. This analysis revealed a significant block × group interaction (*F*(3,117) = 2.77, *P* = 0.045, *η_p_^2^* = 0.066, 95% CI = 0 to 0.149). As training proceeded, participants in the stress group showed a reduced outcome representation at the time of S presentation (first six vs. last six blocks: *t*_22_ = 3.59, *P*_corr_ = 0.004, *d* = 0.748, 95% CI = 0.277 to 1.206), whereas the outcome representation remained rather constant in participants of the control group (first six vs. last six blocks: *t*_17_ = 1.08, *P*_corr_ = 0.590, *d* = 0.255, 95% CI = −0.219 to 0.721, Figure 4). In the last six blocks of the task, the classification accuracy for the outcome was significantly lower in the stress group compared to the control group (*t*_39_ = 3.13, *P*_corr_ = 0.012, *d* = 0.986, 95% CI = 0.326 to 1.635; lower training intensity (first 18 blocks of the task): all *t*_39_ < 1.13, all *P*_corr_ = 1, all *d* < 0.355, all 95% CI = −0.914 to 0.975; note that the overall pattern remains when trials are blocked differently, see Supplementary Information). Strikingly, the reduced outcome representation was significantly correlated with the reduced behavioural sensitivity to the outcome devaluation during Dev O^high^ blocks (Spearman’s ρ = – 0.482, 95% CI = −0.688 to −0.205, *P* = 0.001, Figure 5).

**Figure 4.**
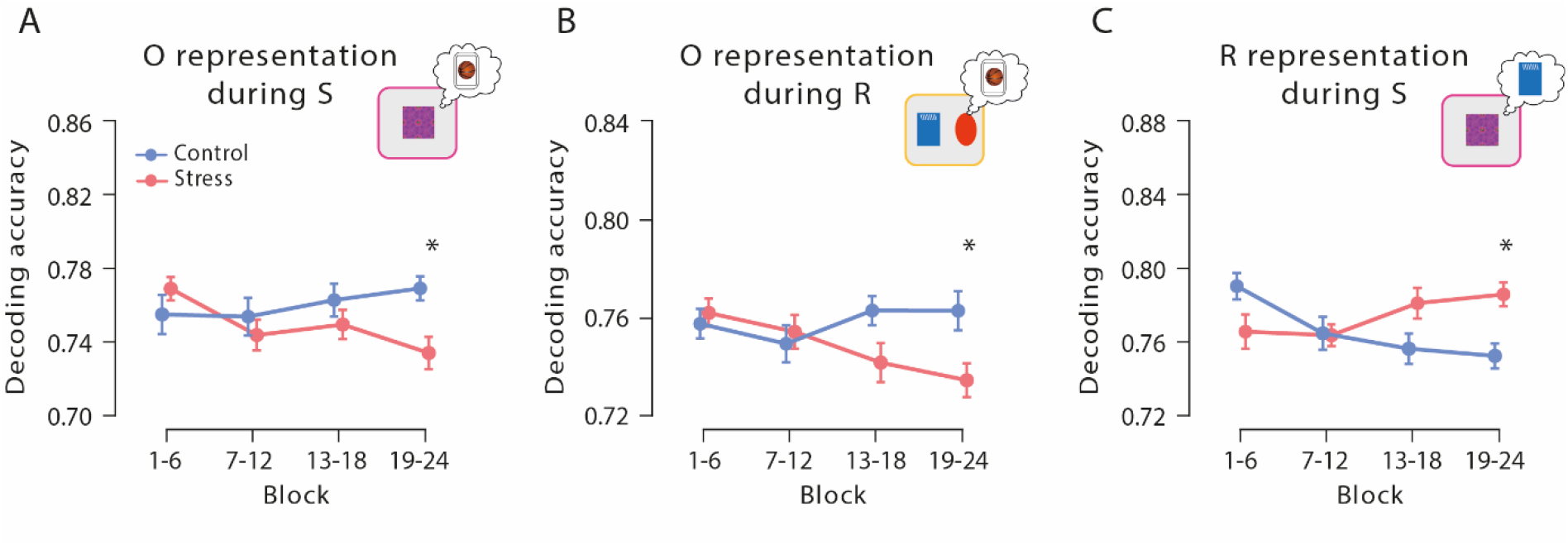
Outcome and response representations throughout the reinforcement learning task. **A** and **B** Outcome representation during stimulus presentation and response choice, respectively. As training proceeded, the outcome representations decreased in the stress group, while there were no changes in the control group. At the end of the learning task, outcome representations were significantly lower in stressed participants than in controls. **C** Response representations during stimulus presentation. Stressed participants showed significantly stronger response representations after extended training compared to the control group. Data represent means ± SEM. * *P* < 0.05, Bonferroni-corrected.

**Figure 5.**
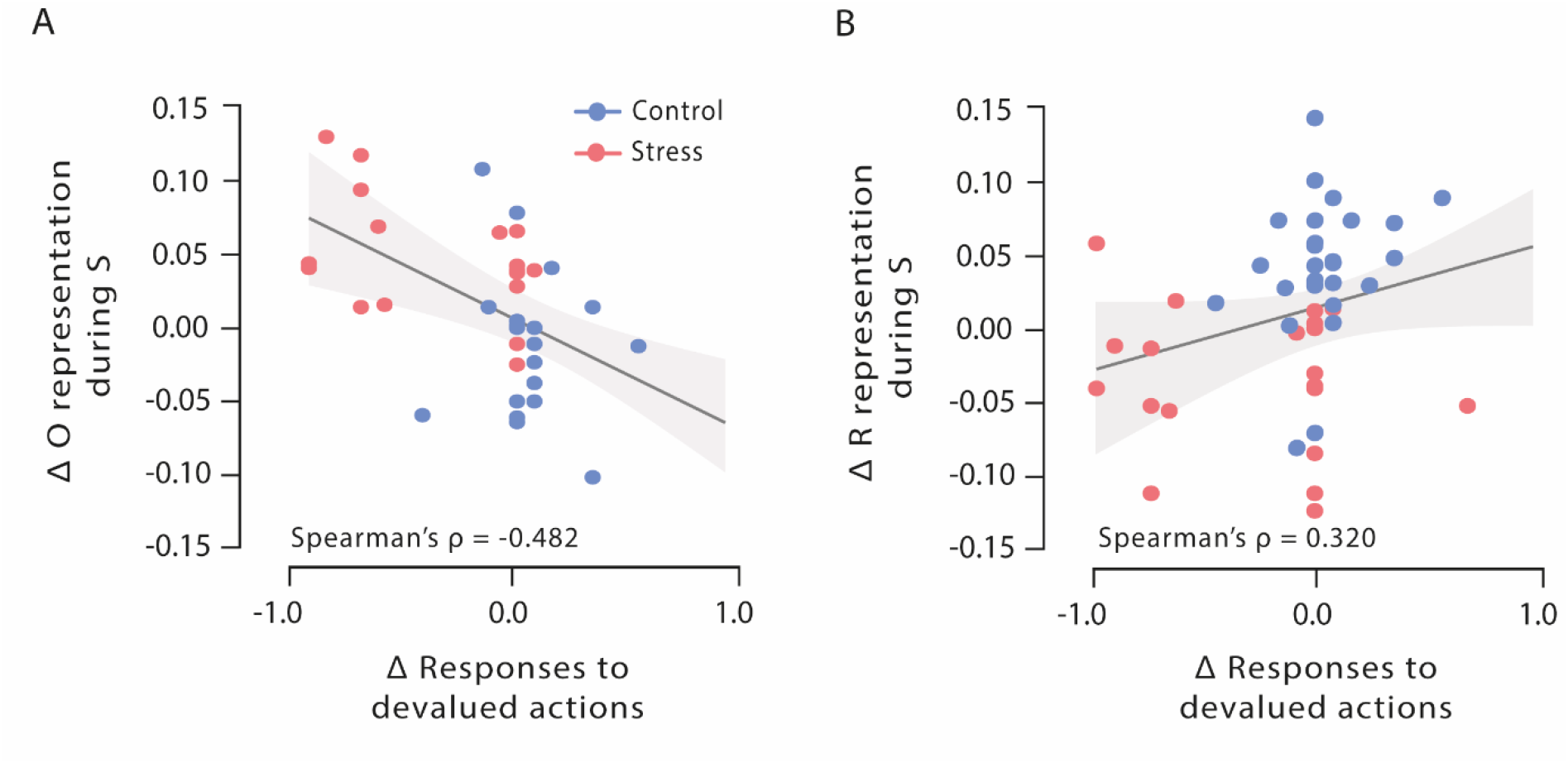
Correlations of outcome and response representation during stimulus presentation with responses to devalued actions during Dev O^high^ blocks. **A** Decrease of outcome representation during stimulus presentation was significantly correlated with the reduced behavioural sensitivity to the outcome devaluation during Dev O^high^ blocks. **B** Increase of response representation was significantly correlated with an increase of responses to devalued actions during Dev O^high^ blocks. Regression lines are added for visualization purpose, the light-coloured background areas indicate its 95% CI intervals.

When we analysed the outcome representations at the time of the choice between the two aliens, a very similar pattern emerged: at the end of the task, participants in the stress group showed a decreased outcome representation at the time point of the choice (*t*_22_ = 2.94, *P* = 0.008, *d* = 0.613, 95% CI = 0.161 to 1.054), whereas there was no such effect in the control group (*t*_17_ = 0.59, *P* = 0.566, *d* = 0.138, 95% CI = −0.328 to 0.600). In addition, stressed participants had reduced outcome representations relative to controls, reflected in a significantly reduced classification accuracy, during the response choice at the end of the reinforcement learning task (block × group interaction: *F*(3,117) = 2.99, *P* = 0.034, *η_p_^2^* = 0.071, 95% CI = 0 to 0.156; stress vs. control, high training intensity: *t*_39_ = 2.75, *P*_corr_ = 0.036, *d* = 0.865, 95% CI = 0.214 to 1.506; lower training intensities: all *t*_39_ < 2.30, all *P*_corr_ > 0.108, all *d* < 0.725, all 95% CI = −0.500 to 1.358). Together, these results show that acute stress reduced, after extended training, the representation of action outcomes which are a hallmark of goal-directed control.

### Stress boosts response representations after extended training

While it is assumed that the outcome representation that is crucial for goal-directed S-R-O learning is reduced with increasing habitual behaviour control, it can be predicted that response (R) representations at the time of stimulus (S) presentation should be strengthened once habitual S-R learning governs behaviour. Therefore, we trained another classifier to discriminate between categories that were used during the reinforcement learning task as response options (red vs. blue alien). This classifier was used to examine changes in response representation during the stimulus presentation (S) throughout the reinforcement learning task. For these response representations, a block × group ANOVA revealed a significant interaction effect (*F*(3,147) = 5.82, *P* < 0.001, *η_p_^2^* = 0.106, 95% CI = 0.021 to 0.192). As shown in Figure 4, participants of the stress group showed a stronger response representation, reflected in a higher classification accuracy for the response categories, with increasing training intensity (first half vs. last half: *t*_25_ = 2.51, *P*_corr_ = 0.038, *d* = 0.491, 95% CI = 0.079 to 0.894), whereas there was even a decrease in the control group (first half vs. last half: *t*_24_ = 3.50, *P*_corr_ = 0.004, *d* = 0.701, 95% CI = 0.256 to 1.134). In the last six blocks of the reinforcement learning task, i.e. after extended training, stressed participants had significantly higher response representations than participants in the control group (*t*_49_ = 2.75, *P*_corr_ = 0.032, *d* = 0.770, 95% CI =0.197 to 1.336; lower training intensities: all *t*_49_ < 1.92, all *P*_corr_ > 0.244, all *d* < 0.537, all 95% CI 0.025 to 1.094). Interestingly, this increase of response representation was significantly correlated with an increase of responses to devalued actions during Dev O^high^ blocks (Spearman’s ρ = 0.320, 95% CI = 0.049 to 0.547, *P* = 0.022, Figure 5). Thus, our MVPA results indicate that stress leads to an increased response representation at the time of stimulus presentation, in line with the predicted increase in habitual S-R learning.

Importantly, when we grouped the classification data not in four blocks consisting of 144 trials in total (averaged over six successive blocks containing 24 reinforcement learning trials each) but in 2, 3, 6 or 12 blocks, the pattern of results for the neural outcome and response representations was largely comparable (Supplementary Table 1).

### Outcome and response representations are uncorrelated

In order to test whether the observed opposite changes in outcome and response representations after stress reflected independent or linked neural representations, we analysed Bayesian correlation between the classification accuracies. These analyses revealed moderate evidence for the null hypothesis that outcome representations, both at stimulus presentation and response selection, were uncorrelated with response representation at choice (both Pearson’s | *r*| < 0.165, both 95% CI = 0.154 to 0.442, both BF01 > 3.092, Figure 6), suggesting that outcome representations and response representations may be independent of each other.

**Figure 6.**
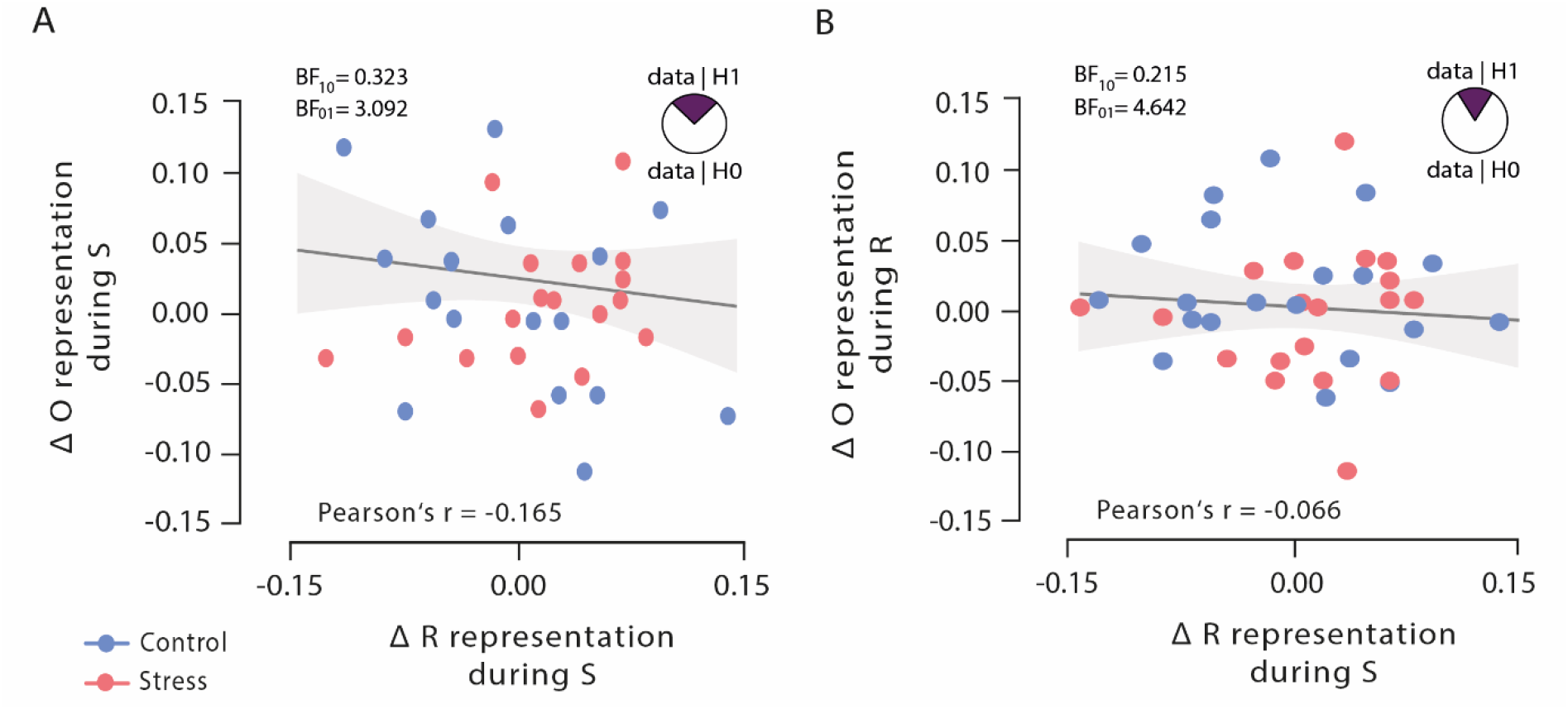
Bayesian correlations of outcome representation during stimulus presentation and response selection with response representation during stimulus presentation. **A** Outcome representation during stimulus presentation was not correlated with response representation during response selection. As visualized in the pie chart, the corresponding Bayes factor suggests that the observed data are 3.092 times more likely under the null hypothesis (H0) than under the alternative hypothesis (H1). **B** Outcome representation during response selection was not correlated with response representation during stimulus presentation. As visualized in the pie chart, the corresponding Bayes factor suggests that the observed data are 4.642 times more likely under the H0 than under the H1. Regression lines are added for visualization purpose, the light-coloured background areas indicate its 95% CI intervals.

### Control variables and performance in the DMS task

At the beginning of the experiment, stress and control groups did not differ in subjective mood (all *t*_56_ < 1.20, all *P* > 0.235, all *d* < 0.316, all 95% CI = −0.326 to 0.833), subjective chronic stress (all *t*_54_ < 1.07, all *P* > 0.290, all *d* < 0.285, all 95% CI = −0.677 to 0.811), depressive mood (*t*_56_ = 1.07, *P* = 0.289, *d* = 0.281, 95% CI = −0.238 to 0.797), state or trait anxiety (both *t*_56_ < 0.44, all *P* > 0.663, both *d* < 0.115, both 95% CI = 0.401 to 0.630, Supplementary Table 1). Behavioural performance in the DMS task, used to train the classifier, was, as expected, very high (average performance: 97.5% correct, *SD* = 0.054) and comparable between groups (*t*_56_ = 0.23, *P* = 0.818, *d* = 0.061, 95% CI = −0.455 to 0.576). The average classification accuracy of the classifier was 72% (*SD* = 0.068) for the response categories (blue rectangular vs. red oval symbol) and 66% (*SD* = 0.046) for the outcome categories (object vs. scene image) and did not differ between the stress and control groups (both *t*_51_ < 0.89, both *P* > 0.376, both *d* < 0.246, both 95% CI = −0.669 to 0.786).

## Discussion

Previous research showed that stress favours habitual responding over goal-directed action *(**Braun and Hauber, 2013; Dias-Ferreira et al., 2009; Gourley et al., 2012; Schwabe et al., 2009; Schwabe et al., 2010; Schwabe et al., 2011b; Schwabe et al., 2012; Seehagen et al., 2015; Smeets et al., 2019; Smeets and Quaedflieg, 2019; Soares et al., 2012; Wirz et al., 2018**).* Although this stress-induced bias towards habits has important implications for stress-related mental disorders *(**Adams et al., 2018; Goeders, 2004; Schwabe et al., 2011a**),* a fundamental question has remained elusive: Is the shift towards habits under stress due to diminished goal-directed or enhanced habitual control (or both)? Canonical behavioural assays of the mode of behavioural control can hardly distinguish between these alternatives. Here, we leveraged EEG-based decoding of outcome and response representations – the key components of goal-directed and habitual processing, respectively – to provide evidence that acute stress results in a decrease of goal-directed processing that is paralleled by an increase in habitual processing.

Our behavioural and ERP data corroborate previous reports of a stress-induced shift from goal-directed to habitual control *(**Braun and Hauber, 2013; Schwabe et al., 2011b; Smeets and Quaedflieg, 2019**).* Specifically, stressed participants showed, after extended training, increased responding to a devalued action, suggesting a reduced behavioural sensitivity to the outcome devaluation which indicates less goal-directed behaviour and more habitual responding *(**Adams and Dickinson, 1981**).* Recent evidence showed that goal-directed and habitual processing may also be reflected in ERPs that are either sensitive or insensitive to changes in the outcome of an action *(**Luque et al., 2017**).* We observed here that stress appears to reduce the sensitivity of late potentials to the outcome devaluation, whereas stress left an occipital P1 component unaffected that was insensitive to the outcome devaluation and may thus be considered habitual *(**Luque et al., 2017**).* However, similar to the behavioural insensitivity to an outcome devaluation after stress, these ERPs cannot separate reduced goal-directed from increased habitual responding. To disentangle goal-directed and habitual processing, we used an MVPA-based decoding approach. We reasoned that since the goal-directed system acquires S-R-O associations whereas the habit system learns S-R associations *(**Adams, 1982; Balleine and Dickinson, 1998; Dickinson, 1985; Dickinson and Balleine, 1994**),* neural representations of outcome O and response R during learning may shed light on specific contributions of the goal-directed and habitual system, respectively. Outcome representations are a key feature of goal-directed learning. However, similar to behavioural measures and ERPs, changes in outcome representations alone are not sufficient to distinguish between enhanced habitual and reduced goal-directed control because both of these alternatives may be reflected in reduced outcome representations. Therefore, we also decoded response R representations which should increase if the habit system takes over but be less sensitive to changes in the goal-directed system. Critically, our Bayesian analyses indicated that changes in outcome and response representations were uncorrelated, which may be taken as evidence that they reflect at least partly dissociable signatures of goal-directed and habitual processing, respectively.

We show that stress led to a reduction in outcome representations, both at the time of stimulus presentation and at the time of action selection, and a parallel increase of response representations during stimulus presentation indicative of enhanced S-R associations (Figure 4). Both the stress-induced reduction in outcome representations and the increase in response representations were directly correlated with the behavioural (in)sensitivity to the outcome devaluation (Figure 5). However, while outcome representations were negatively correlated with participants’ responding to the devalued action, there was a positive correlation for response representations, which might lend further support to the view that these representations reflect distinct processes. Together, our results indicate that acute stress leads to enhanced habitual and impaired goal-directed processing. The latter is in line with evidence showing reduced orbitofrontal activity in the face of elevated stress hormones (***Schwabe et al., 2012***), which is accompanied by increased habitual processing. Based on previous pharmacological studies *(**Gourley et al., 2012; Schwabe et al., 2010; Schwabe et al., 2011b;Schwabe et al., 2012**),* we assume that these opposite effects of stress on goal-directed and habitual processing are based on the concerted action of glucocorticoids and noradrenaline on the neural substrates of goal-directed and habitual control.

It might be argued that an alternative way of disentangling goal-directed and habitual contributions to instrumental responding under stress would be through an analysis of modelbased and model-free learning. Model-based learning uses representations of the environment, expectations, and prospective calculations to predict future value. Model-free learning, in turn, acquires action-values directly based on past experiences, without building an internal model of the environment (***Daw et al., 2005; Dayan and Berridge, 2014***). It has been argued that model-based learning enables goal-directed action, whereas model-free learning underlies habitual responding *(**Dolan and Dayan, 2013**).* In line with a stress-induced shift towards habits, stress appears to impair model-based learning (***Otto et al., 2013; Radenbach et al., 2015***). However, although there are relevant parallels between modelbased vs. model-free learning on the one hand and goal-directed vs. habitual behaviour on the other hand, the relation between these processes is not necessarily clear-cut *(**Dezfouli and Balleine, 2013**)* and experimental evidence for a direct link between model-based/model-free learning and goal-directed/habitual behaviour is scarce. Moreover, there are important differences between these concepts. For instance, while model-free learning can be present after only few trials, establishing habit behaviour requires extensive training *(**Adams, 1982; Dickinson et al., 1995; Tricomi et al., 2009; Wit et al., 2018**).* Furthermore, while model-free learning is based on previously experienced outcomes, habit learning is defined as S-R learning without links to the outcome engendered by a response *(**Adams, 1982; Adams and Dickinson, 1981; Balleine and Dickinson, 1998**).* Thus, we assume that an analysis of model-based and model-free learning cannot fully capture the separate contributions of the goal-directed and habit systems to instrumental behaviour under stress and that the present decoding of neural representations of outcome and response representations goes significantly beyond what could be expected from an analysis of model-based and model-free computations.

Importantly, both our behavioural and our neural decoding data showed that stress affected the balance of goal-directed and habitual processes only after extended training. Previous human studies could not distinguish between such early and late effects of stress because the mode of behavioural control was only assessed at the end of training *(**Schwabe and Wolf, 2009; Smeets et al., 2019**).* The present results, however, dovetail with rodent data showing that (chronic) stress effects on the control of instrumental learning occurred only after extended training *(**Dias-Ferreira et al., 2009**).* In general, it is commonly assumed that early training is under goal-directed control while extensive training results in habitual control(***Adams, 1982; Dickinson et al., 1995; Tricomi et al., 2009***). Whether overtraining may induce habits in humans is currently debated (***Wit et al., 2018***) and our data cannot speak to this issue as training may have been too limited to result in overtraining-related habits. Findings on ‘cognitive’ and ‘habitual’ forms of navigational learning in rats demonstrated that stress hormones may accelerate a shift from ‘cognitive’ to ‘habit’ learning that would otherwise only occur after extended training *(**Packard, 1999**).* It is tempting to speculate that a similar mechanism might be at work during instrumental learning in stressed humans. Future studies are required to test this hypothesis by using the present neural decoding approach in combination with an extensive training protocol and groups of participants that are exposed to stress at distinct stages of the learning process.

Goal-directed action and habits are commonly considered to be two sides of the same coin. If a behaviour is more goal-directed, it is automatically less habitual (and vice versa). Obviously, a behaviour cannot be fully goal-directed and habitual at the same time, according to canonical operational definitions *(**Adams, 1982; Dickinson and Balleine, 1994**).* However, behaviour may not always necessarily be either fully goal-directed or habitual and there may be different degrees to which behaviour is under goal-directed or habitual control. In particular, given that the two modes of behavioural control are subserved by distinct neural circuits (***Balleine and O’Doherty, 2010***), it should be possible to separate goal-directed and habitual contributions to learning at a specific point in time. Classical behavioural paradigms can hardly disentangle goal-directed and habitual components in a specific pattern of responding (e.g. insensitivity to outcome devaluation). Furthermore, tests of related cognitive functions, such as inhibitory control, provide only indirect evidence, if any, on the balance of goal-directed and habitual processes. We therefore used here a MVPA-based decoding approach that focussed on neural representations that are a hallmark feature of goal-directed and habitual processing, respectively. Using this novel approach, we show that acute stress reduces outcome representations and, at the same time, increases response representations in healthy humans, suggesting that stress exerts opposite effects on goal-directed and habitual processing that manifest in the predominance of habitual behaviour under stress.

## Methods

### Participants and design

Sixty-two healthy volunteers participated in this experiment. This sample size was based on earlier studies on stress and mnemonic control in our lab *(**Schwabe and Wolf, 2012**)* and a-priori power calculation using G*POWER 3 suggesting that this sample size would be sufficient to reveal a medium-sized effect in a mixed-design ANOVA with a power of .80. Exclusion criteria were checked in a standardized interview before participation and comprised any current or chronic mental or physical disorders, medication intake or drug abuse. Further, smokers and women taking hormonal contraceptives were excluded from participation because previous studies revealed that smoking and hormonal contraceptive intake may alter the cortisol response to stress. In addition, participants were asked to refrain from food intake, caffeine, and physical activity for 2 hours before testing. Four participants had to be excluded from analysis due to medication intake shortly before participation, thus leaving a final sample of 58 participants (32 men, 26 women; age: *M* = 29.53 years, *SEM* = 2.57 years). Participants received a monetary compensation of 30€ for study participation, plus a performance-dependent compensation (2 to 5€). All participants gave written informed consent before entering the study, which was approved by the local ethics committee. In a between-subjects design, participants were randomly assigned to the stress (15 men, 15 women) or control group (17 men, 11 women).

### Stress and control manipulation

Participants in the stress condition underwent the Trier Social Stress Test (TSST; ***Kirschbaum et al., 1993***), a standardized stress protocol known to reliably elicit both subjective and physiological stress responses *(**Allen et al., 2014; Kirschbaum et al., 1993***). Briefly, the TSST consisted of a mock job interview during which participants were asked to give a 5 minute free speech about why they are the ideal candidate for a job tailored to their interests and a 5 minute mental arithmetic task (counting backwards in steps of 17 from 2043 as fast and accurate as possible; upon a mistake, they had to stop and start again from 2023). Both, the free speech and the mental arithmetic task were performed in front of a rather cold and non-reinforcing panel of two experimenters (1 man, 1 woman) that were dressed in white coats and introduced as experts in ‘behavioural analysis’. Furthermore, participants were videotaped throughout the TSST and could see themselves on a large screen placed next to the panel. In the control condition, participants gave a 5 minute speech about a topic of their choice (e.g. their last holiday) and performed a simple mental arithmetic task (counting in steps of two) for 5 minutes while being alone in the experimental room; no video recordings were taken. During the control condition, the experimenter waited in front of the door outside the room where he/she was able to hear whether the participants complied with the instructions.

To assess the effectiveness of the stress manipulation, subjective and physiological measurements were taken at several time points across the experiment. More specifically, participants rated the stressfulness, difficulty, and unpleasantness of the previous experience immediately after the TSST or control manipulation on a scale from 0 (“not at all”) to 100 (“very much”). In addition, blood pressure and pulse were measured using an OMRON M400 device (OMRON, Inc., MI) before, during, immediately after the TSST/control manipulation and 20, 60 and 105 minutes after the TSST/control manipulation. To quantify cortisol concentrations and elevations during the experiment, saliva samples were collected from participants using Salivette^®^ collection devices (Sarstedt, Germany) before, during, as well as 20, 60 and 105 minutes after the TSST/control manipulation. Saliva samples were stored at – 20°C until the end of testing. At the end of data collection, we determined the free fraction of cortisol from the saliva samples using a commercially available luminescence assay (IBL, Germany).

### Reinforcement Learning Task

In order to investigate goal-directed and habitual contributions to behaviour, we used a modification of a recently introduced reinforcement learning task *(**Luque et al., 2017**).* In this reinforcement learning task, participants played the role of space traders on a mission to trade fractal stimuli (either a pink or a green fractal) for playing cards (either scene or object playing cards) with aliens from two tribes (“red alien tribe” and a “blue alien tribe”). Each tribe traded only one type of playing cards (i.e. either scene or object playing cards). One type of cards (the high-value outcome, O^high^, worth 100 points) was more valuable than the other one (the low-value outcome, O^low^, worth 20 points). In addition, one tribe of aliens exchanged a playing card only for a specific fractal (e.g. the red alien tribe only traded scene playing cards and only for a pink fractal but not for a green fractal). If a fractal was given to the alien tribe, the exchange was rejected by the alien (e.g. if the participant wanted to trade the green fractal with the red alien). In that case, participants did not receive a playing card and hence did not gain any points. Importantly, participants had to pay 5 points for every successful exchange (response costs). If a playing card was given to the incorrect alien and the exchange was denied, however, there were no response costs. Participants were encouraged to earn as many points as possible. At the end of the experiment, participants received a monetary reward dependent on the number of points earned throughout the reinforcement task.

Because the critical difference between goal-directed action and habitual responding is whether behaviour is sensitive to an outcome devaluation or not *(**Adams and Dickinson, 1981**),* specific outcomes were devalued in some of the task blocks. Participants were told that both types of cards needed to be kept in different boxes (scene cards in a “scene box” and objects cards in an “object box”) but that sometimes one box was full and had to be emptied. In that case, participants were not able to store cards of the respective category anymore. Thus, in these blocks, the cards were worthless regardless of their usual value (i.e. the outcome was devaluated).

The status of the boxes (available/not available) varied only between blocks. Participants were informed about the box status via a message on the screen at the beginning of each block. In blocks without outcome devaluation *(NoDev),* both types of cards could be kept in their respective boxes and had their usual value. In blocks with devaluation of the high-valued outcome (*Dev O^high^),* O^high^ cards could not be stored and thus their new value was zero, whereas O^low^ cards could be stored and thus had still the usual value. In blocks with devaluation of the low-valued outcome (*Dev O^low^),* O^low^ cards could not be saved and had no value anymore, while O^high^ cards still had high value (Figure 1). Importantly, due to the response costs (5 points) for each card exchange, giving a card to an alien that traded the devalued card category was detrimental and should be avoided if behaviour is under goal-directed control.

Reinforcement learning trials were composed of stimulus-response-outcome (S-R-O) sequences. At the beginning of each trial, a fixation cross was presented in the centre of the computer screen for a random duration between 800 to 1200 ms. Then, one of the two highly distinct fractals (high-value stimulus, S^high^, or low-value stimulus, S^low^) was presented. After 2000 ms, pictures of two aliens were shown at either side of the screen. The two aliens represented two different response options (R1 and R2). Participants were instructed to choose via button press to which alien they wanted to give the fractal. After their response, the screen was cleared and the outcome feedback was appeared on the screen for 1000 ms. If participants chose the incorrect alien, the message “Error!” was presented. If they did not respond within 2 seconds, the message “Time out, please respond faster!” was displayed. The specific stimuli used as S^high^ and S^low^, R1 and R2, and O^high^ and O^low^, respectively, were counterbalanced across participants. The position of the aliens (left/right) representing the response options was defined randomly for each participant at the beginning of the experiment.

The feedback provided on each trial varied between blocks. During NoDev blocks, participants saw the value of the card that was gained before as well as the corresponding response costs (“+100/-5” for O^high^ and “+20/-5” for O^low^). In Dev blocks (Dev O^high^ or Dev O^low^ blocks), this information was masked for all trials. Before Dev blocks, participants were informed that the outcome feedback was unavailable during these blocks because of a solar interference (in devaluation blocks, participants saw on the last screen of each trial “???” instead of value), but that this solar interference would not influence the value of the cards that they could gain.

In addition to reinforcement learning trials, we included consumption trials in each block to test whether participants understood the task structure. At the beginning of the experiment, participants were instructed that sometimes the aliens become distracted and participants could take one of the two types of cards without trading it for fractals and therefore without paying response costs. In these consumption trials, the message “The aliens seem distracted …” was shown for 1000 ms, followed by a countdown (from 3 to 1) in the center of the screen (over 3 s). Then, one scene and one object card appeared, one on the left and the other on the right (positions were selected randomly for each consumption trial). Participants choose one card via button press and their choice revealed whether they were aware of the value of the respective stimuli in the respective block (NoDev, Dev O^high^, Dev O^low^). Outcome screens were identical to those of the reinforcement learning trials.

Participants completed 27 NoDev blocks, 3 Dev O^high^ blocks, and 3 Dev O^low^ blocks, with 27 trials each: 12 learning trials with S^high^, 12 learning trials with S^low^, and three consumption trials. Reinforcement learning trials within each block were presented randomly. Consumption trials were displayed at trial numbers 7, 14, and 21 within each block. The O^high^ was always devaluated during the 2^nd^, 16^th^ and 29^th^ block (Dev O^high^), whereas O^low^ was always devalued in the 3^rd^, 17^th^ and 30^th^ block (Dev O^low^). Thus, these Dev blocks were presented after initial, moderate, and extensive training, respectively.

At the end of each block, participants saw how many cards they had gained in the previous block, the total point value of the cards earned in that block, and the total number of points they had gained until then. Then, the next block started with the information about which box(es) was/were available (i.e., whether any outcome was devalued) for the upcoming block.

Performance during the devalued trials revealed the degree to which instrumental behaviour was goal-directed or habitual. Goal-directed action is indicated by the formation of S-R-O associations. Thus, if participants used a goal-directed system, they should adapt their responses to the actual outcome value in case of outcome devaluation. During devalued trials, participants did not earn any points if they choose the response associated with the devalued outcome. Importantly, however, they had to pay response costs leading to a subtraction of points. Thus, goal-directed participants should choose the response option associated with a trade rejection. In that case, participants did not have to pay any response costs (Figure 1). In contrast, habitual behaviour is reflected in simpler S-R associations rendering instrumental behaviour insensitive to changes in outcome value. Hence, choosing the action associated with the devalued outcome (albeit no points could be earned and response costs had to be paid) indicated less goal-directed and more habitual behaviour.

### Delayed-matching-to-sample task

In order to analyse the neural representations of response options and action outcomes, we trained an EEG-based classifier (see below) on a delayed-matching-to-sample (DMS) task. This task was presented before and after participants had completed the reinforcement learning task to avoid any time-dependent biases in the trained classifier. In each DMS task, participants completed 128 trials. Participants were presented four different target types: object cards, scene cards, blue and red symbols. In addition to colour, symbols also differed in shape, line orientation and line position (blue rectangles with left-oriented lines in the upper area vs. red ovals with right-oriented lines in the lower area). Pictures used as targets were selected randomly from a pool of 256 pictures (90 × object cards, 90 × scene cards, 48 × blue symbols and 48 × red symbols) with the restriction that successive trials did not belong to the same category more than three times in a row. The remaining pictures were used as targets during the second DMS task. On each trial, the target was shown for 2 seconds at the centre of a computer screen. Participants were asked to keep it in mind during a subsequent delay phase of 2 seconds during which they saw a blank screen. Then, a probe stimulus was presented (the target and a distractor belonging to the same category) and the participants had to indicate via button press which picture they saw before. The position of the target during the response choice (right vs. left) was randomized. Different stimuli were used in the second DMS task compared to the first one.

### Control variables

In order to control for potential group differences in depressive mood, chronic stress, and anxiety, participants completed the Beck Depression Inventory (BDI; ***Beck et al., 1996***), the Trier Inventory for the Assessment of Chronic Stress (TICS; ***Schulz and Schlotz, 1999***), and State-Trait Anxiety Inventory (STAI; ***Spielberger, 1983***) at the end of the experiment. In addition, participants completed a German mood questionnaire (MDBF; ***Steyer et al., 1994***) that measures subjective feeling on three dimensions (elevated vs. depressed mood, wakefulness vs. sleepiness, and calmness vs. restlessness) at the beginning of the experiment.

### Procedure

All testing took place in the afternoon between 1 pm and 8 pm. Participants were instructed to refrain from excessive exercise, and food or caffeine intake in the two hours before testing. After participants’ arrival at the laboratory, EEG was prepared, blood pressure measurements were taken and a first saliva sample was collected. Moreover, participants completed the mood scale *(**Steyer et al., 1994**).* Then, participants performed the first DMS task. After completing this task, participants received written instructions about the reinforcement learning task. In order to further familiarize participants with the structure of this task, participants completed a 5-minute short tutorial afterwards. Next, participants underwent either the TSST or the control manipulation. Immediately thereafter, subjective assessments of this manipulation and another saliva sample were collected and blood pressure was measured again. Next, participants were briefly reminded of the instructions for the reinforcement learning task they had received before. Twenty minutes after the TSST/control manipulation, when cortisol was expected to have reached peak levels (***Kirschbaum et al., 1993***), participants collected another saliva sample before they started with the reinforcement learning task. After the 15^th^ reinforcement task block and after finishing the reinforcement task, further saliva samples were collected and blood pressure was measured again (~ 60 minutes and ~ 105 minutes after stress onset). Finally, participants performed the second DMS task.

### Statistical analysis

Subjective and physiological stress responses were analysed by mixed-design ANOVAs with the within-subject factor time point of measurement and the between-subject factor group (stress vs. control). Participants’ responses in the reinforcement learning task were subjected to mixed-design ANOVAs with the within-subject factor stimulus type (S^high^ and S^low^), outcome devaluation (noDev: 1^st^, 12^th^ and 28^th^ block; Dev O^high^: 2^nd^, 16^th^ and 29^th^ block; Dev O^low^: 3^rd^, 17^th^ and 30^th^ block), time point (first, second, third) and the between-subject factor group. Significant interaction effects were followed by appropriate post hoc tests. All reported *p* values are two-tailed and were Bonferroni corrected (*P*_corr_) when indicated. Statistical analyses were calculated using SPSS 25 (IBM SPSS Statistics) and JASP version 0.13.0.0 software (www.jasp-stats.org). For the correlations between outcome representations and response representation, we performed Bayesian analysis using JASP version 0.13 software (https:/jasp-stats.org/) and the default Cauchy prior 0.707. For one participant, we obtained only invalid trials during the last Dev O^low^ block. Thus, this participant could not be included in analyses of Dev O^low^. Furthermore, two participants did not complete the Trier Inventory for the Assessment of Chronic Stress *(**Schulz and Schlotz, 1999**).* Data of 5 participants had to be excluded from the EEG analysis because of technical failure during the EEG, leaving a sample of 53 participants (control: *n* = 25; stress: *n* = 28) for EEG analyses.

### EEG recordings

During the DMS and reinforcement learning task, participants were seated approximately 80 cm from the computer screen in an electrically shielded and sound attenuated cabin. EEG was recorded using a 128-channel BioSemi ActiveTwo system (BioSemi, Amsterdam, The Netherlands) organized according to the 10-5 system digitized at 2024 Hz. Additional electrodes were placed at the left and right mastoids, approximately 1 cm above and below the orbital ridge of each eye and at the outer canthi of the eyes. The EEG data were online referenced to the BioSemi CMS-DRL (common mode sense-driven right leg) reference. Electrode impedances were kept below 30 kΩ. EEG was amplified with a low cut-off frequency of 0.53 Hz (= 0.3 s time constant).

### EEG analysis

#### Preprocessing

Preprocessing was performed offline using FieldTrip *(**Oostenveld et al., 2011**)* and EEGLAB *(**Delorme and Makeig, 2004**)* as well as custom scripts implemented and processed in MATLAB (The Mathworks, Natick, MA). The PREP pipeline procedure (***Bigdely-Shamlo et al., 2015***) was utilized to transform the channel EEG data using a robust average reference. In addition, channels were interpolated using the *spherical* option of EEGLAB *eeg_interp* function. Then, data were filtered with a high pass filter of 0.1 Hz and a low pass filter of 100 Hz and and downsampled to 250 Hz. For MVPA, epochs from the DMS task (2000 ms relative to the delay onset) and from the reinforcement learning task (2000 ms relative to the onset of stimulus and response option presentation, respectively) were extracted. For ERP analysis, EEG data were segmented into epochs from −200 to 2000 ms around the stimulus onset and baseline-corrected by subtracting the average 200 ms prestimulus interval.

#### ERP analysis

Based on previous studies *(**Hickey et al., 2010; MacLean and Giesbrecht, 2015**),* we expected to find an effect of stimulus value (i.e. the difference between activity elicited by S^high^ and S^low^) in the occipital P1 component, peaking within the time window from 75 to 200 ms relative to stimulus onset. Because all stimuli were presented centrally, P1 activity was analysed in the midline occipital electrode Oz. The P1 peak was defined as the largest positive peak between 75 and 200 ms after the stimulus onset at Oz (averaging across all conditions). A time window of 70 ms around that peak was then selected for analysis, in line with previous research *(**Luque et al., 2017**).* Because the P1 maximum amplitude was at 115 ms from stimulus onset across participants, the P1 magnitude for each condition was defined as the mean EEG signal across the 80 to 150 ms time window.

Based on previous research on the neural underpinnings of goal-directed action *(**Luque et al., 2017**),* we further assessed effects of stress on brain activity over centro-parietal regions (CPz) at a later time window. To this end, ERP data from 400 to 700 ms was subdivided into six consecutive, non-overlapping time bins with a duration of 50 ms each. The later ERP data were analysed by mixed-design ANOVAs with the within-subject factors outcome devaluation (NoDev, Dev O^high^ and Dev O^low^) and stimulus value (S^high^ and S^low^) and the between-subject factor group. Significant outcome devaluation × stimulus value × group interaction effects were appropriately corrected for multiple comparisons.

#### MVPA training

The multivariate decoding analyses were implemented using the MVPA-Light toolbox (***Treder, 2020***). The classifier was trained within-subject using a linear support vector machine (SVM) on the preprocessed data of the DMS task (delay phase). All EEG channels were used as features. To improve the signal to noise ratio for MVPA, trials were averaged to pseudo trials *(**Isik et al., 2014**).* Each pseudo trial was an average of two trials. We performed two separate analyses corresponding to the following different classes: object vs. scene (SVM^object/scene^) and blue symbol vs. red symbol (SVM^blue/red^). To identify the optimal time window for decoding per participant, we implemented a sliding window averaging 100 ms with a step size of 10 ms. The SVM^object/scene^ and SVM^blue/red^ with the highest performance were used to decode the neural outcome and response representation, respectively. Generalization of the classifier was evaluated using a leave-one-out procedure. If the classifier’s performance is significantly above chance, it indicates that the EEG patterns contain class-specific information and that the class can be reliably decoded from the EEG data (***Murphy et al., 2011***). The chance level in a simple 2-class paradigm is not exactly 50% but 50% with a confidence interval at a certain alpha level. Therefore, we calculated this interval utilizing the Wald interval with adjustments for a small sample size *(**Agresti and Caffo, 2000; Müller-Putz et al., 2008**).* The threshold for chance performance was 63.59% for the classification of blue vs. red symbols and 60.11% for object vs. scene images. Participants with classification accuracy below chance in the DMS task were not included in subsequent analyses. During the classification of blue vs. red symbols, the highest performance of two participants did not exceed the threshold for chance performance. During the classification of blue vs. red scenes, classification accuracy of 13 participants was not significant. Hence, the sample for analysis of the outcome representation (based on SVM^object/scene^) and response representation (based on SVM^red/blue^) reported in the main text was 41 (control: n = 18; stress: n = 23) and 51 (control: n = 25; stress: n = 26) participants, respectively. Importantly, however, we computed all analyses again including all participants regardless of significant classification scores. This additional analysis left our findings largely unchanged (see Supplementary Results). Furthermore, the classification accuracy of the first and the second DMS task did not differ (blue/red symbol classification: *F*(1,51) = 0.66, *P* = 0.798, *ƞp^2^* = 0.013, 95% CI = 0 to 0.127; DMS session × group interaction: *F*(1,51) = 0.03, *P* = 0.863, *rp* = 0.001, 95% CI = 0 to 0.024; object/scene classification: *F*(1,51) = 2.87, *P* = 0.096, *ƞp^2^* = 0.053, 95% CI = 0 to 0.203; DMS session × group: *F*(1,51) = 0.03, *P* = 0.860, *ƞp^2^* = 0.006, 95% CI = 0.003 to 0.011). Thus, we pooled the data from both DMS task sessions and trained an overall classifier to ensure that the classifier was not affected by any time-related biases and that there is a sufficient number of trials to train a reliable classifier.

#### MVPA decoding

The SVM^object/scene^ and SVM^blue/red^ trained on the independent DMS dataset were used to assess the outcome (object card vs. scene card) and response (blue alien vs. red alien) representation, respectively, in the trial-by-trial reinforcement learning task during the NoDev blocks. Both classifiers were applied to the respective test data using an overlapping sliding window with a time average of 100 ms and a step size of 10 ms. The maximal classification accuracy indicated the strength of outcome and response representation.

Goal-directed behaviour is characterized by an action-outcome (S-R-O) association, which should result in an anticipatory outcome representation during stimulus presentation and response choice. Hence, we applied the SVM^object/scene^ to the stimulus presentation and response choice phase of the reinforcement learning task in order to decode the outcome representation. With increasing habitual behaviour control of behaviour, outcome representations should be reduced, while response representations should be even increased due to the formation of S-R associations. Therefore, we applied the SVM^blue/red^ to the stimulus presentation phase to decode the response representation. Maximal accuracy values were then averaged over six blocks to get a reliable indicator of classification accuracy. Statistical analyses were performed at the group level averaging across individual decoding accuracies. Decoding accuracies were analysed by mixed-design ANOVAs with the within-subject factor block (1-6, 7-12, 13-18 and 19-24 block) and the between-subject factor group. In addition, we computed Spearman correlations between Δ classification scores (averaged classification accuracy during the first six blocks minus the classification accuracy during the last six blocks) and Δ responses to devalued action for O^high^ blocks (responses to devalued action during the first devaluation block, i.e. the 2^nd^ overall block, minus responses to devalued action during the last devaluation block, i.e. the 29^th^ overall block).

## Acknowledgements

We thank Keyvan Khatiri, Charlotte Germer, Marian Wiskow and Yichen Zhong for their assistance during data collection.

## Competing interests

The authors declare no competing interests.

## Additional information

### Funding

Universität Hamburg.

### Author contributions

Jacqueline K Meier, Data curation, Formal analysis, Investigation, Visualization, Methodology, Writing—original draft, Project administration, Writing—review and editing; Bernhard P Staresina, Methodology, Writing—review and editing; Lars Schwabe, Conceptualization, Supervision, Methodology, Writing—original draft, Writing—review and editing, Funding acquisition.

### Ethics

Human subjects: University of Hamburg approved the study. All participants gave informed consent to participate in the EEG study.

## Additional files

### Data availability

Data reported in this manuscript are available from the website: https://github.com/08122019/From-goal-directed-action-to-habit.

## SUPPLEMENTAL MATERIAL

### Supplementary Results

**Supplementary Figure 1.**
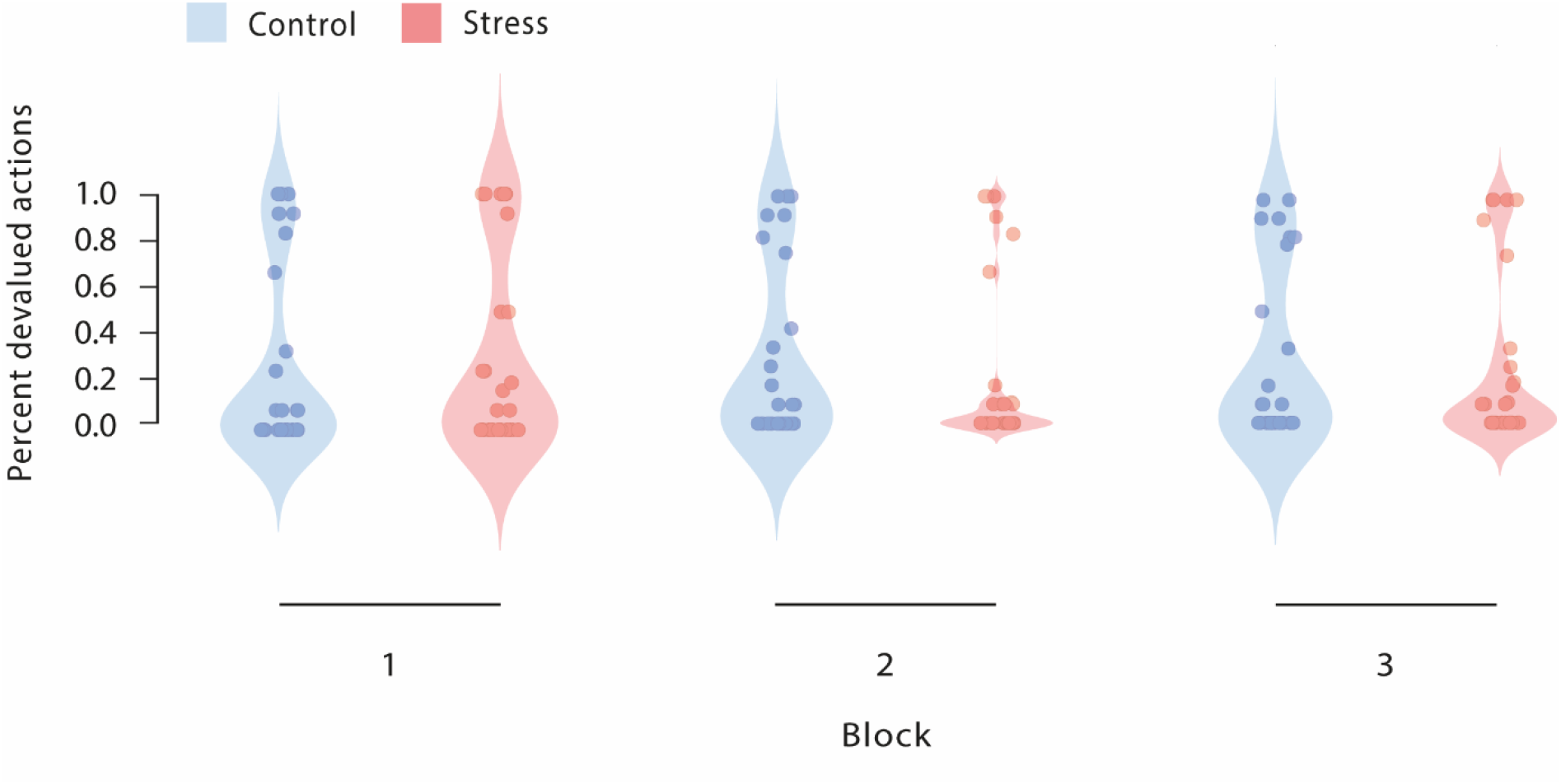
Percentage of responses to devalued actions across the reinforcement learning task during Dev O^low^ blocks.

**Supplementary Figure 2.**
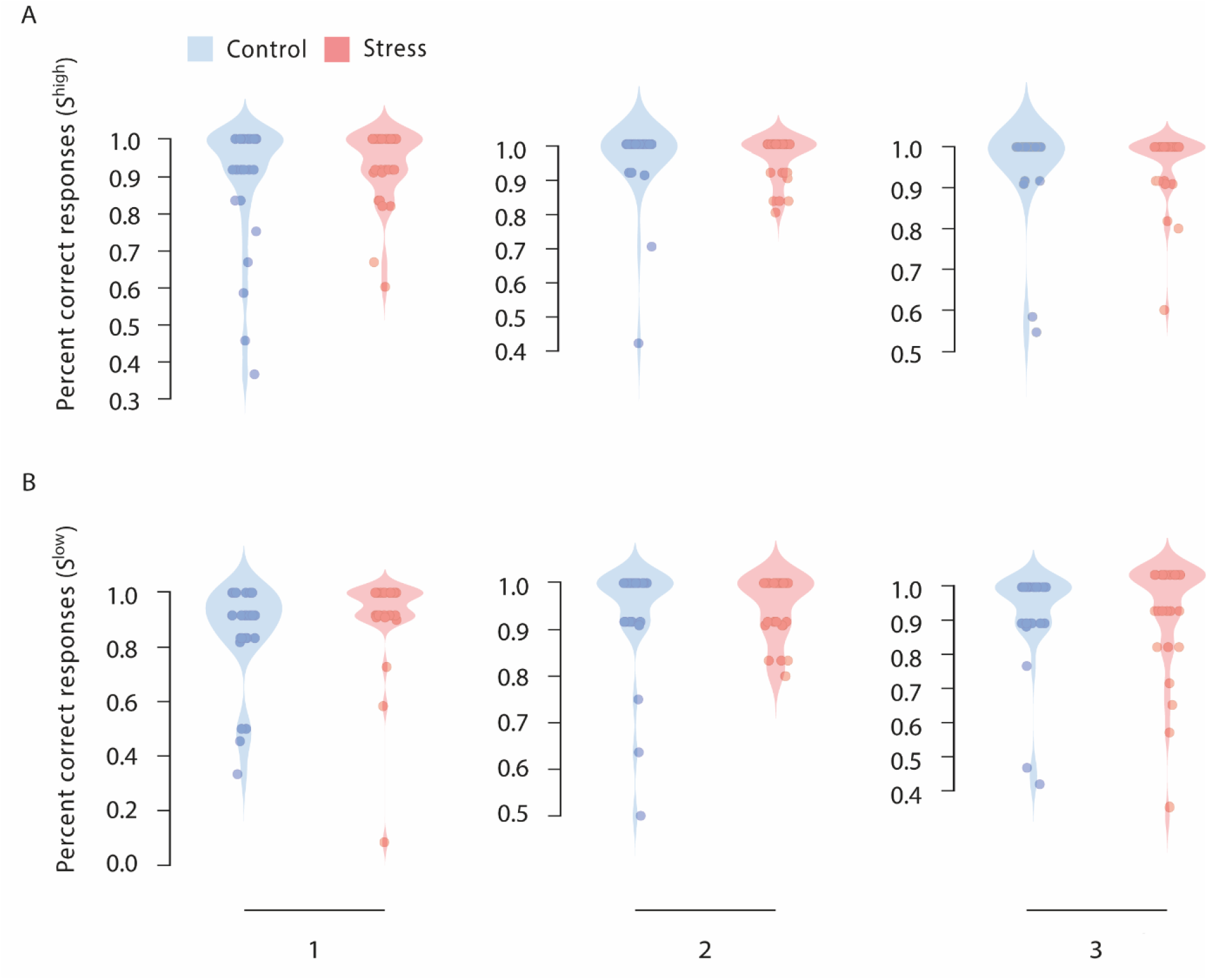
Percentage of correct responses during NoDev blocks. **A** Percentage of correct responses to S^high^ during NoDev blocks. **B** Percentage of correct responses to S^low^ during NoDev blocks.

### Stress alters event-related potentials associated with goal-directed and habitual processing

Although our analyses of the neural data focussed mainly on the decoding of outcome and response representations, there is recent evidence suggesting that habitual and goal-directed processes might also be reflected in event-related potentials (ERPs) *(**Luque et al., 2017**).* The extent to which a reward related ERP is sensitive to an outcome devaluation or not is assumed to indicate the degree of habitual or goal-directed processing. Our data show that the occipital stimulus-locked P1 component was insensitive to outcome devaluation (outcome devaluation × stimulus value interaction: *F*(2, 102) = 0.63, *P* = 0.536, *η_p_^2^* = 0.012, 95% CI = 0 to 0.069, Supplementary Figures 3–5) suggesting the formation of an “attentional habit” *(**Luque et al., 2017**).* This component was not modulated by stress (stimulus value × group: *F*(1,51) = 0.13, *P* = 0.723, *η_p_^2^* = 0.002, 95% CI 0 to 0.086; outcome devaluation × stimulus value × group: *F*(2,102) = 0.11, *P* = 0.900, *η_p_^2^* = 0.002, 95% CI = 0 to 0.028). Moreover, we identified a late component that tended to be sensitive to the outcome devaluation during Dev O^high^ blocks in control participants (devalued vs. valued: *t*_24_ = 1.91, *P* = 0.068, *d* = 0.382, 95% CI = −0.028 to 0.785) but not in stressed participants (devalued vs. valued: *t*_27_ = 1.57, *P* = 0.127, *d* = 0.297, 95% CI = −0.084 to 0.673; outcome devaluation × stimulus value × group interaction: *F*(2,102) = 5.2, *P*_corr_ = 0.042, *η_p_^2^* = 0.093, 95% CI = 0.008 to 0.199; stimulus value × group interaction: *F*(1,51) = 6.05, *P* = 0.017, *η_p_^2^* = 0.106, 95% CI = 0.003 to 0.273; no such effect in NoDev and Dev O^low^ blocks: stimulus value × group interaction: both *F*(1,51) < 1.44, both *P* > 0.236, both *η_p_^2^* < 0.027, both 95% CI = 0 to 0.159).

**Supplementary Figure 3.**
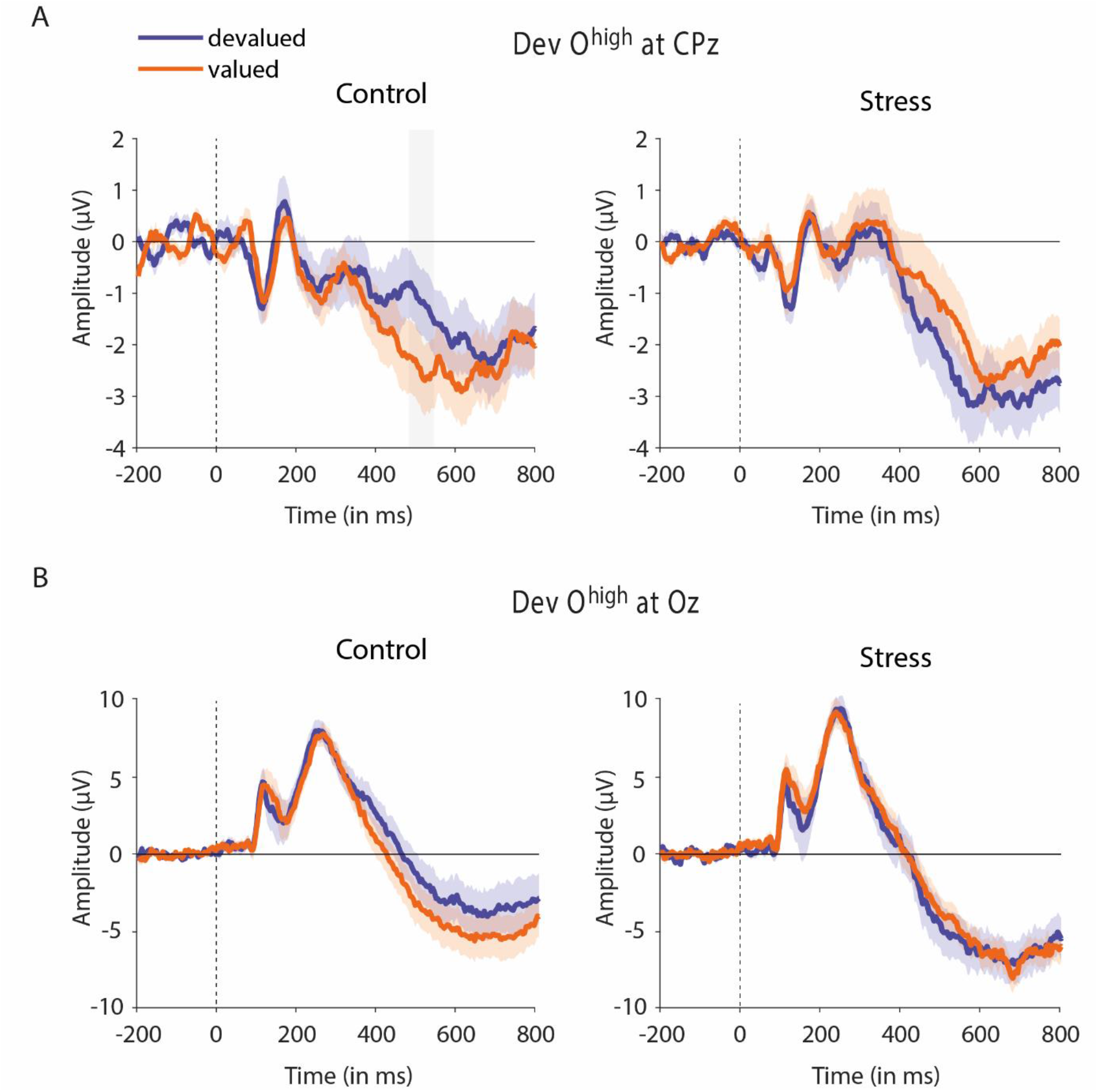
Event-related potentials during Dev O^high^ blocks. **A** Mean centroparietal activity for devalued and valued stimuli during Dev O^high^ for control and stressed participants (baseline-corrected). The late component tended to be sensitive to the outcome devaluation in the control group but not in the stress group. The light-colored background bar refers to the time range showing marginally significant group differences. **B** Mean occipital activity for devalued and valued stimuli during Dev O^high^ for control and stressed participants (baseline-corrected). The stimulus-locked P1 component was insensitive to outcome devaluation in both groups. Data represent means ± SEM.

**Supplementary Figure 4.**
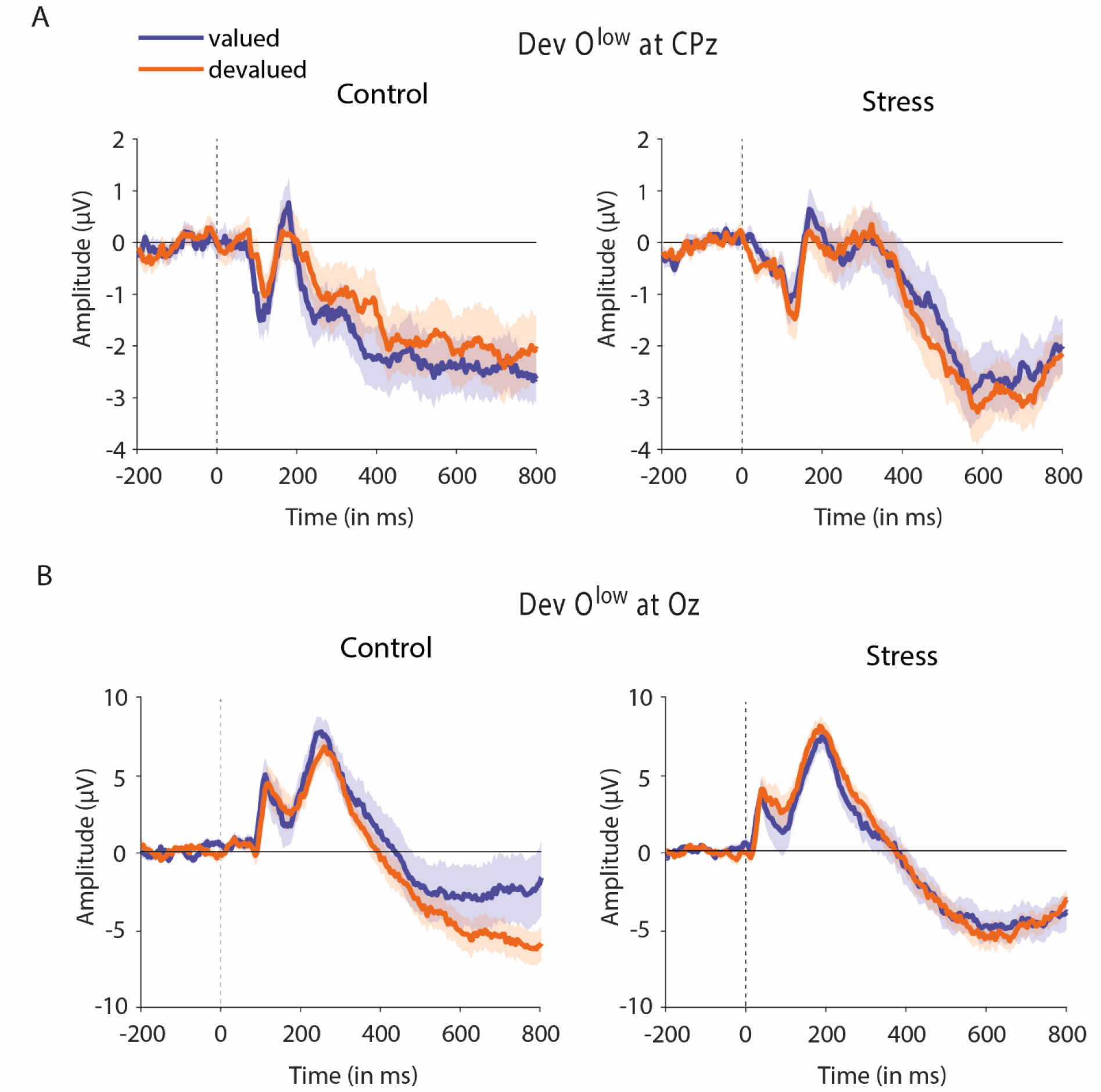
Event-related potentials during Dev O^low^ blocks. **A** Mean centroparietal activity for devalued and valued stimuli during Dev O^low^ for control and stressed participants (baseline-corrected). **B** Mean occipital activity for devalued and valued stimuli during Dev O^low^ for control and stressed participants (baseline-corrected). Data represent means ± SEM.

**Supplementary Figure 5.**
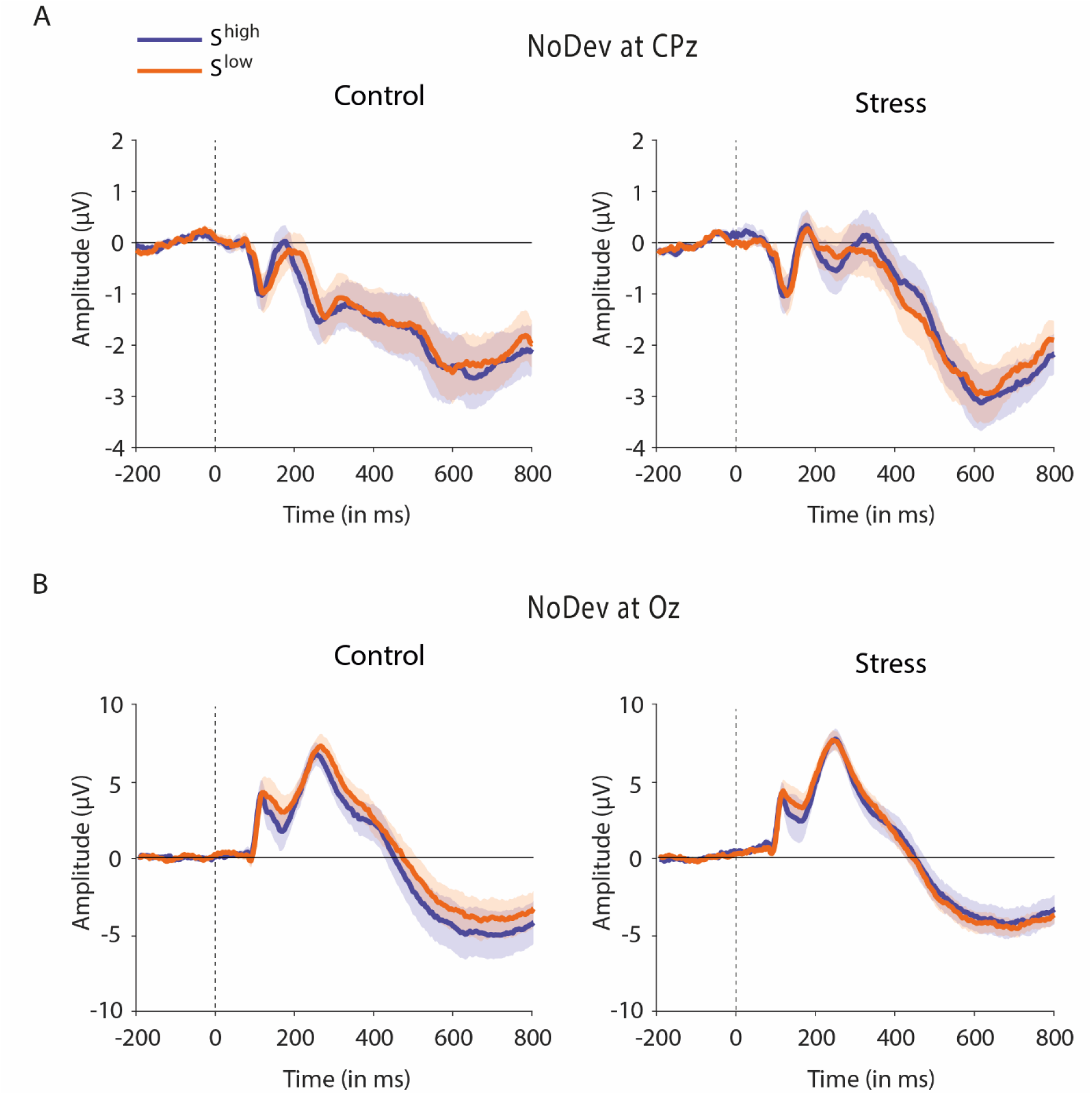
Event-related potentials during NoDev blocks. **A** Mean centroparietal activity for for S^high^ and S^low^ during NoDev for control and stressed participants (baseline-corrected). **B** Mean occipital activity for S^high^ and S^low^ during NoDev for control and stressed participants (baseline-corrected). Data represent means ± SEM.

### Stress effects on outcome and response representations for alternatively grouped data

The overall pattern of stress effects on outcome and response representations largely remains when trials are blocked differently (Supplementary Table 1).

**Supplementary Table 1.**
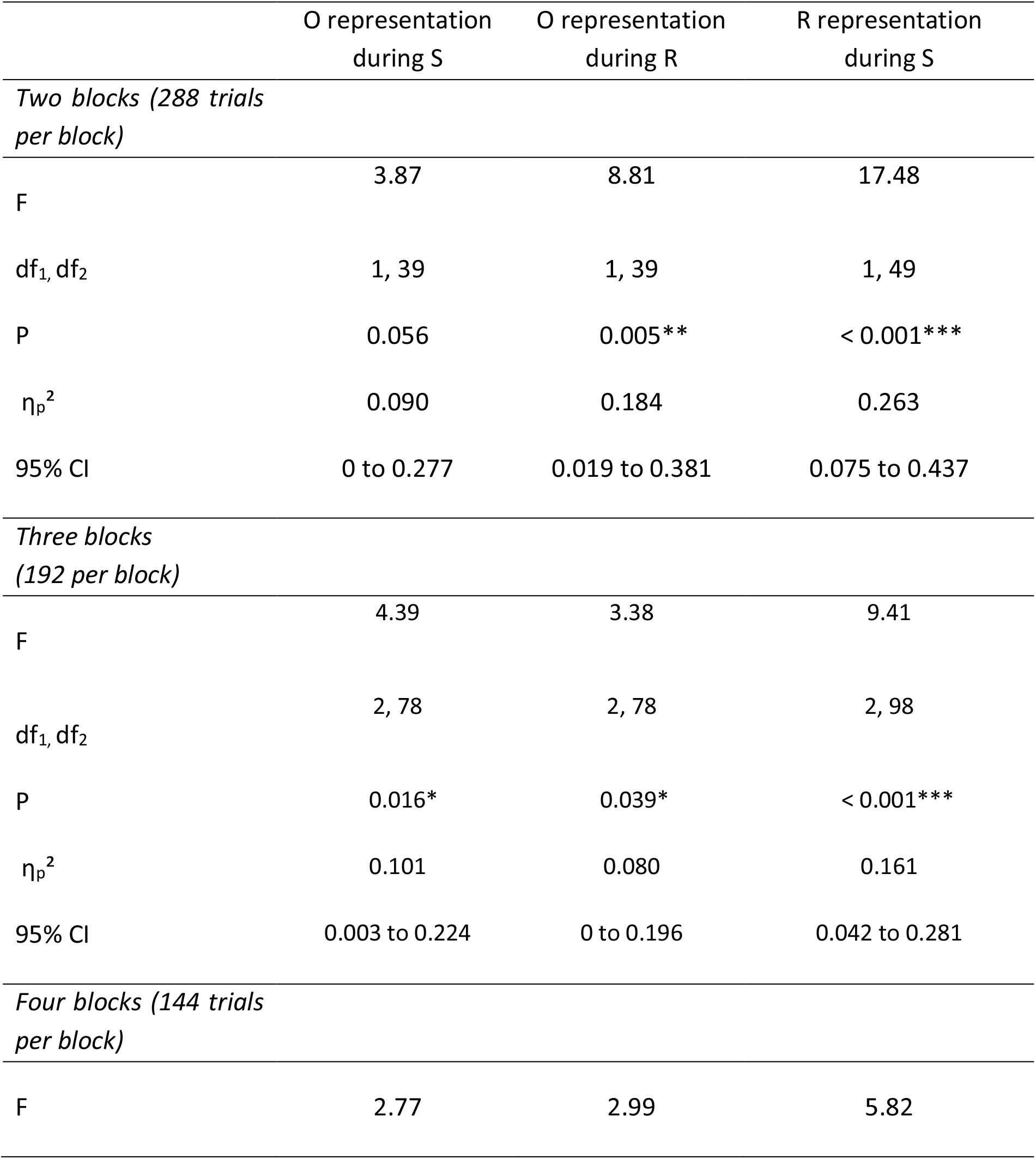
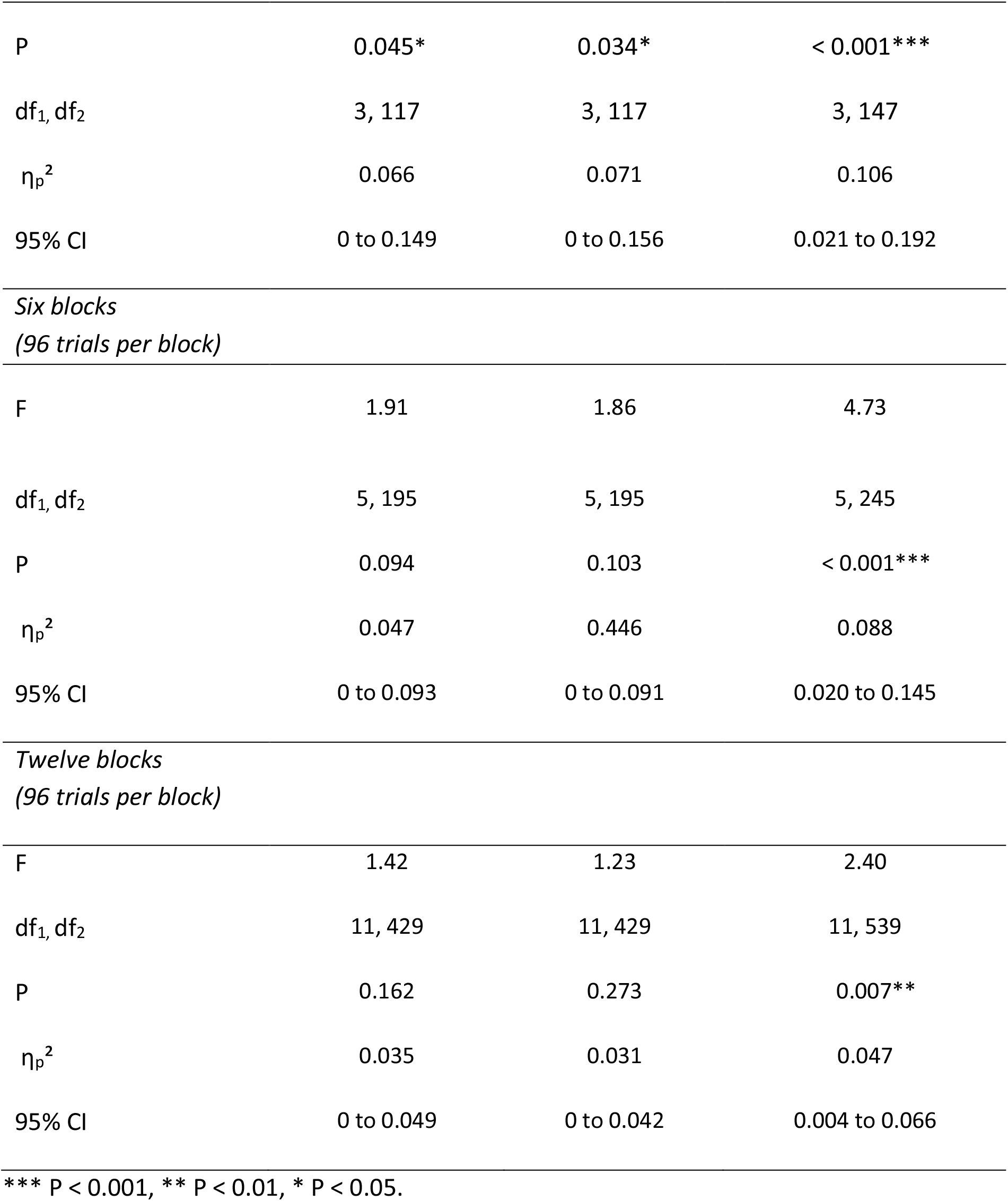
Classification results (block × group interaction) block for alternatively grouped data.

### Stress effects on outcome representations for N = 53 (including those with decoding accuracy below chance in the DMS task)

The analysis of outcome representations during the stimulus presentation revealed a significant block × group interaction (*F*(3,153) = 4.44, *P* = 0.005, *η_p_^2^* = 0.080, 95% CI = 0.008 to 0.158). As training proceeded, participants in the stress group showed a reduced outcome representation at the time of S presentation (first six vs. last six blocks: *t*_27_ = 4.21, *P* < 0.001, *d* = 0.796, 95% CI = 0.365 to 1.216), whereas the outcome representation remained rather constant in participants of the control group (first vs. last block: *t*_24_ = 1.32 *P* = 0.199, *d* = 0.264, 95% CI = −0.660 to 0.138). In the last six blocks of the task, the classification accuracy for the outcome was significantly lower in the stress group compared to the control group (*t*_51_ = 2.95, *P*_corr_ = 0.020, *d* = 0.811, 95% CI = 0.246 to 1.369; lower training intensity (first 18 blocks of the task): all *t*_51_ < 1.03, all *P*_corr_ = 1, all *d* < 0.283, all 95% CI = −0.977 to 0.824). Reduced outcome representation tended to correlate with the reduced behavioural sensitivity to outcome devaluation during Dev O^high^ blocks (Spearman’s ρ = −0.253, 95% CI = −0.490 to 0.019, *P* = 0.068, *n* = 53).

When we analysed the outcome representations at the time of the choice between the two aliens, a similar pattern emerged: at the end of the task, participants in the stress group tend to showed a decreased outcome representation at the time point of the choice (*t*_27_ = 1.66, *P* = 0.109, *d* = 0.313, 95% CI = −0.069 to 0.690), whereas there was no such effect in the control group (first six vs. last six blocks: *t*_24_ = 1.50, *P* = 0.147, *d* = 0.300, 95% CI = 0.104 to 0.698). In addition, stressed participants had reduced outcome representations relative to controls, reflected in a significantly reduced classification accuracy, during the response choice in the last six blocks of the reinforcement learning task (block × group interaction: *F*(3,153) = 3.72, *P* = 0.013, *η_p_^2^* = 0.068, 95% CI = 0.003 to 0.142; stress vs. control, high training intensity: *t*¾1 = 2.69, *P*_corr_ = 0.032, *d* = 0.757, 95% CI = 0.195 to 1.313; lower training intensities: all *t*_51_ < 1.65, all *P*_corr_ > 0.420, all *d* < 0.454, all 95% CI = −0.870 to 0.998). Together, these results show that acute stress reduced, after extended training, the representation of action outcomes which are a hallmark of goal-directed control.

### Stress effects on response representations for N = 53 (including those with decoding accuracy below chance in the DMS task)

While it is assumed that the outcome representation that is crucial for goal-directed S-R-O learning is reduced with increasing habitual behaviour control, it can be predicted that response (R) representations at the time of stimulus (S) presentation should be strengthened once habitual S-R learning governs behaviour. Therefore, we trained another classifier to discriminate between categories that were used during the reinforcement learning task as response options (red vs. blue alien). This classifier was used to examine changes in response representation during the stimulus presentation (S) throughout the reinforcement learning task. A block × group ANOVA revealed a significant interaction effect (*F*(3,153) = 3.96, *P* = 0.009, *η_p_^2^* = 0.072, 95% CI = 0.005 to 0.147). In the last six blocks of the reinforcement learning task, i.e. after extended training, stressed participants tended to have higher response representations than participants in the control group (*t*_51_ = 1.96, *P* = 0.056, *d* = 0.538, 95% CI =-1.085 to 0.014; lower training intensities: all *t*_51_ < 1.74, all *P* > 0.087, all *d* < 0.480, all 95% CI −0.070 to 1.025).

**Supplementary Table 2.** Control variables.

|  | Control |  | Stress |  |
| --- | --- | --- | --- | --- |
|  | M | SEM | M | SEM |
| *MDBF scales* |  |  |  |  |
| Elevated vs. depressed mood | 17.54 | 0.35 | 17.80 | 0.30 |
| Sleepiness vs. wakefulness | 15.43 | 0.61 | 14.43 | 0.56 |
| Calmness vs. restlessness | 17.21 | 0.50 | 16.73 | 0.44 |
| *BDI* | 8.79 | 1.27 | 6.90 | 1.23 |
| *STAI scales* |  |  |  |  |
| State anxiety | 34.07 | 1.41 | 35.07 | 1.76 |
| Trait anxiety | 35.71 | 1.73 | 36.27 | 1.64 |
| *TICS scales* |  |  |  |  |
| Work overload | 18.36 | 1.12 | 18.43 | 1.19 |
| Social overload | 13.61 | 0.95 | 12.29 | 0.79 |
| Performance pressure | 21.64 | 1.41 | 22.68 | 1.13 |
| Work discontent | 18.32 | 0.96 | 17.96 | 1.22 |
| Excessive workload | 11.61 | 0.70 | 12.43 | 0.92 |
| Lack of social recognition | 8.82 | 0.54 | 8.21 | 0.47 |
| Social tension | 10.71 | 0.78 | 11.32 | 0.67 |
| Social isolation | 13.68 | 0.97 | 13.61 | 0.99 |
| Chronic worrying | 8.54 | 0.54 | 8.32 | 0.59 |
| TICS screening scale | 25.29 | 1.49 | 25.07 | 1.50 |

MDBF, Multidimensional Mood Questionnaire; BDI, Beck Depression Inventory;
STAI, State-Trait Anxiety Inventory; TICS, Trier Inventory of Chronic Stress.

## Notes

### Competing Interest Statement

The authors have declared no competing interest.

